# Dissociable representations of decision variables within subdivisions of macaque orbitofrontal and ventrolateral frontal cortex

**DOI:** 10.1101/2024.03.10.584181

**Authors:** Frederic M. Stoll, Peter H. Rudebeck

**Author notes:** **CORRESPONDENCE SHOULD BE ADDRESSED TO:** Dr. Frederic M. Stoll and Dr. Peter H. Rudebeck Icahn School of Medicine at Mount Sinai One Gustave L. Levy Place New York, NY, 10029, USA.

## Abstract

Ventral frontal cortex (VFC) in macaques is involved in many affective and cognitive processes and has a key role in flexibly guiding reward-based decision-making. VFC is composed of a set of anatomically distinct subdivisions that are within the orbitofrontal cortex, ventrolateral prefrontal cortex, and anterior insula. In part, because prior studies have lacked the resolution to test for differences, it is unclear if neural representations related to decision-making are dissociable across these subdivisions. Here we recorded the activity of thousands of neurons within eight anatomically defined subregions of VFC in macaque monkeys performing a two-choice probabilistic task for different fruit juices outcomes. We found substantial variation in the encoding of decision variables across these eight subdivisions. Notably, ventrolateral subdivision 12l was unique relative to the other areas that we recorded from as the activity of single neurons integrated multiple attributes when monkeys evaluated the different choice options. Activity within 12o, by contrast, more closely represented reward probability and whether reward was received on a given trial. Orbitofrontal area 11m/l contained more specific representations of the quality of the outcome that could be earned later on. We also found that reward delivery encoding was highly distributed across all VFC subregions, while the properties of the reward, such as its flavor, were more strongly represented in areas 11m/l and 13m. Taken together, our work reveals the diversity of encoding within the various anatomically distinct subdivisions of VFC in primates.

**SIGNIFICANCE STATEMENT:** Ventral frontal cortex (VFC) is essential for flexible decision-making and is composed of many anatomically defined subdivisions. How neural representations related to decision-making vary or not between these subdivisions is unclear. Here we recorded single neuron activity from eight anatomically distinct subdivisions of VFC while macaques made choices between stimuli based on the probability of receiving different flavored fruit juices. We report that neural representations across these subdivisions were dissociable. Area 12l exhibiting the most integrated representations of decision variables at the level of single neurons. By contrast, activity in area 12o was closely related to reward probability whereas activity in area 11m/l and 13m represented juice flavor. Thus, neural representations are distinct across anatomically separable parts of VFC.

## INTRODUCTION

Ventral frontal cortex in primates is involved in a multitude of cognitive and affective functions and plays a prominent role in reward-guided decision-making (Rushworth et al., 2011; Padoa-Schioppa and Conen, 2017; Murray and Rudebeck, 2018). Specifically, lesions or focal disruption of activity in either of two part of the ventral frontal cortex, orbitofrontal cortex (OFC) or ventrolateral prefrontal cortex (vlPFC) impact the ability of humans and non-human primates to choose adaptively based on different costs and benefits (Izquierdo and Murray, 2004; Rudebeck et al., 2013b, 2017; Murray et al., 2015; Reber et al., 2017; Folloni et al., 2021). Similarly, neural activity within OFC and vlPFC encodes the amount, effort, delay, risk, probability, and other option attributes or costs that have to be considered when making a choice (Tremblay and Schultz, 1999; Padoa-Schioppa and Assad, 2006; Kennerley and Wallis, 2009; Kobayashi et al., 2010; Chau et al., 2015; Kaskan et al., 2017; Stoll and Rudebeck, 2023). Furthermore, the level of encoding of decision attributes differs in OFC and vlPFC (Rich and Wallis, 2014; Stoll and Rudebeck, 2023) and this difference corresponds to the dissociable effects of lesions of these two areas (Rudebeck et al., 2017).

OFC and vlPFC are, however, not anatomically homogeneous structures. Within these parts of the frontal lobe, researchers have identified subdivisions that are distinct based on their cytoarchitecture and receptor densities (Walker, 1940; Morecraft et al., 1992; Carmichael and Price, 1994, 1995a, 1995b, 1996; Ongur and Price, 2000). For instance, area 12 is composed of four unique areas that are dissociable based on different anatomical features such as the presence of a granule cell layer, degree of laminar SMI-32 staining, or dopamine receptor densities to name just a few (Carmichael and Price, 1994; Rapan et al., 2023). Subdivisions of OFC and vlPFC are also distinct based on the connections that they send and receive from other parts of the prefrontal cortex, temporal, premotor, subcortical, and sensory areas (Carmichael and Price, 1995a, 1995b, 1996). Such distinct connectivity between dissociable subdivisions of ventral frontal is also observable at the level of fMRI functional connectivity in both human and monkeys (Kahnt et al., 2012; Rapan et al., 2023). Despite these clear differences, it is less clear if such anatomical distinctions are associated with variation in function across these subdivisions of OFC and vlPFC.

A number of studies have revealed both antero-posterior and medial-lateral functional differences within ventral frontal cortex (Kobayashi et al., 2010; Klein-Flügge et al., 2013; Rich and Wallis, 2014; Murray et al., 2015). For instance, Murray and colleagues found that anterior OFC area 11 and posterior OFC area 13 make dissociable contributions to updating and using the current value of rewards to guide choices in a reinforcer devaluation task (Murray et al., 2015). By contrast, Rich & Wallis (2014) reported that there was a gradient of encoding within the medio-lateral extent of ventral frontal cortex, where a greater proportion of neurons within lateral regions represented the valence of outcome that macaque would receive for choosing a course of action. Extending this work to directly link the patterns of activity to anatomically distinct subdivisions of ventral frontal cortex would provide further insight into variation in function in ventral frontal cortex.

Here, we recorded the activity of thousands of single neurons from ventral frontal cortex of macaque monkeys while they performed a two-choice probabilistic task for different outcome flavors. During and following recordings, we confirmed the location of electrodes using CT scans and post-mortem histological reconstruction. This latter approach allowed us to identify eight cytoarchitectonic subdivisions within OFC, vlPFC and AI. When we compared representations across these subdivisions, we found marked differences between neural representations. Notably, neurons in area 12l exhibited the most integrated representations of decision variables whereas neural activity within area 12o was closely related to reward probability. OFC areas 11m/l and 13m exhibited differential representations of outcome flavor over time.

## METHODS

### Subjects

Subjects consisted in two adult male rhesus macaques (*Macaca mulatta*), monkeys M and X. They were 8 and 5.5 years old, and weighed 11.9 and 7.9 kg, respectively, at the start of the neurophysiological recordings. Animals were grouped-housed, kept on a 12-h light dark cycle and had access to food 24 hours a day. Throughout training and testing each monkey’s access to water was controlled for 5 days per week. All procedures were reviewed and approved by the Icahn School of Medicine Animal Care and Use Committee.

### Apparatus

Monkeys were trained to sit head restrained in a custom primate chair situated 56 cm from a 19-inch monitor screen. Choices were reported using gaze location, which was monitored and acquired at 90 frames per second using an infrared oculometer (PC-60, Arrington Research, Scottsdale, AZ). Juice rewards were delivered to the monkey’s mouth using custom-made air-pressured juice dispenser systems (Mitz, 2005). Trial events, reward delivery and timings were controlled using MonkeyLogic behavioral control system (https://monkeylogic.nimh.nih.gov), running in MATLAB (version 2014b, The MathWorks Inc.). Raw electrophysiological activity was recorded using an Omniplex data acquisition system (Plexon, Dallas, TX) and sampled at a 40kHz resolution. Spikes from putative single neurons were automatically clustered offline using the MountainSort plugin of MountainLab (Chung et al., 2017) and later curated manually based on the principal component analysis, interspike interval distributions, visually differentiated waveforms, and objective cluster measures (Isolation > 0.75, Noise overlap < 0.2, Peak signal to noise ratio > 0.5, Firing Rate > 0.05 Hz) (Stoll and Rudebeck, 2023).

### Behavioral task

Monkeys were trained to perform 3 closely related tasks during each session, including a Single option, Instrumental and Dynamic probabilistic tasks (Stoll and Rudebeck, 2023). Our current analyses focused on instrumental trials only. In this task, monkeys could choose between two options presented simultaneously on the right and left side of the screen. Each option was composed of two features: an external-colored rectangle indicating which outcome flavor (out of 2 possible juices per session) monkeys could earn and a second central rectangle, more or less filled, indicating the probability at which this particular outcome flavor would be delivered at the end of the trial. During a given session, monkeys were faced with options containing one of two possible colors (randomly selected from a set of 9 colors) associated with two different juice flavors (randomly picked from a set of 5, which included apple, cranberry, grape, pineapple and orange juices, diluted in 50% water). Probabilities used were from 10% to 90% (by steps of 20% for monkey M and 10% for monkey X).

Monkeys initiated a trial by fixating a central fixation cross for 0.7 to 1.3s (steps of 0.3s). Monkeys were then free to look at the two options, each composed of an outcome flavor and probability, which were displayed for 0.4 to 0.8s (steps of 0.2s, pseudorandomly). The stimulus was then turned off for 0.2s and two response boxes appeared on both sides of the previously shown options (3 possible locations equidistant to the options’ locations; bottom left/right, center left/right, top left/right). Monkeys had to fixate the response box on the side of the desired option within 8s to make their choice. To register their choice, subjects had to maintain fixation on the response box corresponding to that option for a minimum of 0.25s, at which time the other response box associated with the unchosen option would disappear. The 0.25s fixation time required to signal their choice meant that subjects could change their choice within a trial. Continued fixation was required for an additional 0.6 to 1.2s (steps of 0.3s). At that point, the selected response boxes would disappear for 0.3 to 0.7s (steps of 0.2s). Feedback was then delivered. Here, both options were presented again at the same locations, with the selected stimulus option initially flashing (5 times 0.1s ON followed by 0.1s OFF, total time of 0.5s, indicated by 5x in **Fig. 1A**), before staying on the screen for the duration of the reward (if delivered) and an additional 0.5s. In rewarded trials, monkeys received 2-3 pulses of 0.03-0.06s of fluid (separated by 0.1s each, 0.25-0.36ml total reward per trial of the outcome flavor and at the probability indicated by the selected option). Nonrewarded trials were matched in time to trials that included reward delivery. Finally, rewarded trials were followed by a 2s intertrial interval (ITI). Unrewarded trials were followed by a 3.5-4s ITI. If monkeys failed to maintain fixation when required, a large red circle was presented at the center of the screen for 1s, followed by a longer intertrial interval (4-6s for monkey M, 3-4s for monkey X). Failure to initiate a trial by looking at the fixation star within 6s of its appearance resulted in the same red circle and associated ITI.

**Figure 1.**
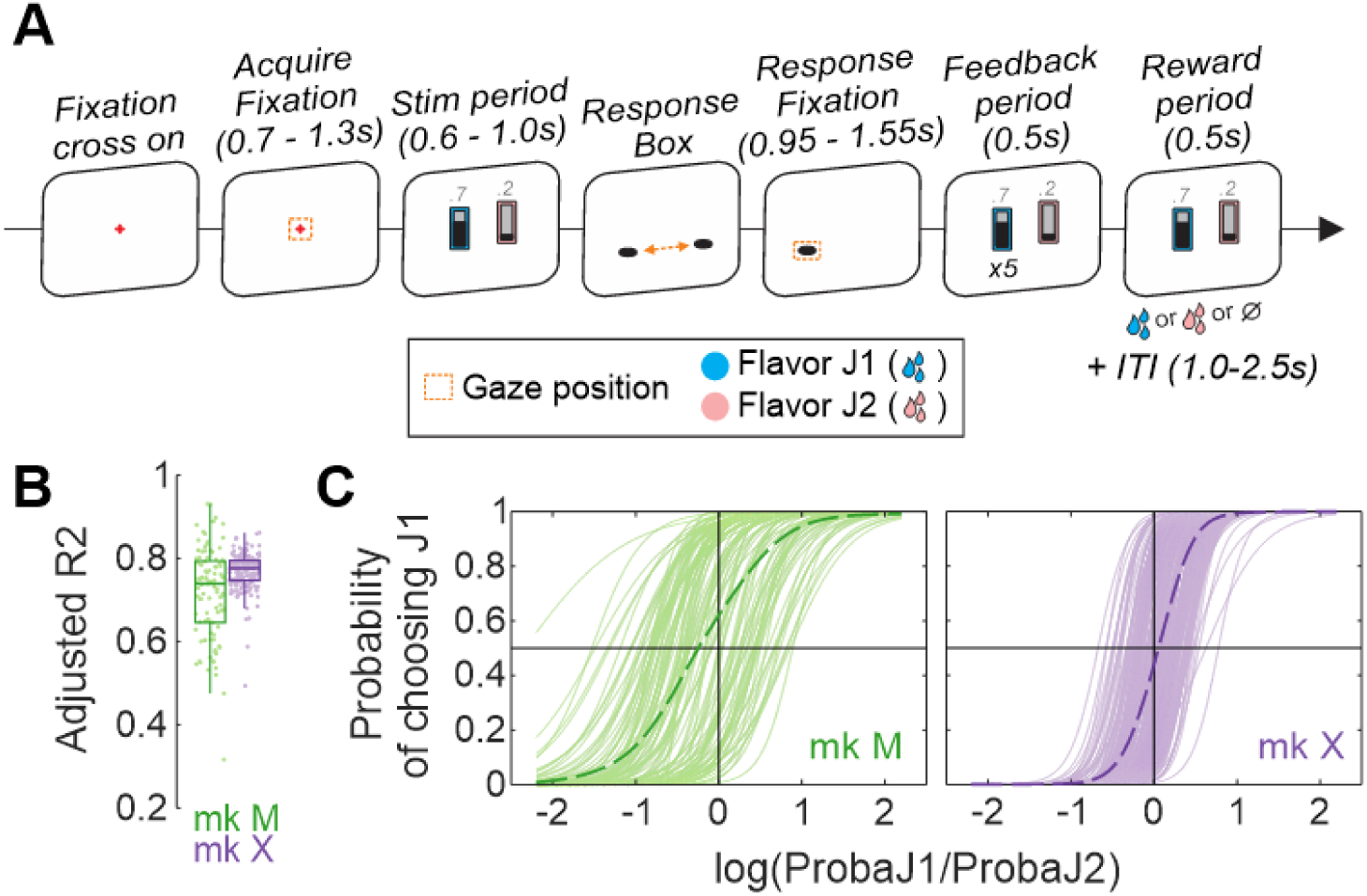
Task and recording locations. (**A**) Schematic representation of the behavioral task. Following a central fixation period, monkeys were shown a set of two stimuli, each comprised of a background color (indicating the outcome flavor that could be earned) and a central gauge (indicating the probability of receiving such outcome). Monkeys reported their choices by maintaining fixation on one of two response boxes located on the side of the chosen stimulus. After a feedback period, a reward was delivered (or omitted) given the characteristics indicated by the chosen stimulus (flavor and probability). (**B**) Adjusted coefficient of determination observed when monkeys’ choices were fitted using a logistic regression that included the log ratio of probabilities associated with each outcome flavor and the flavor of the chosen outcome on the previous trial (*Prev Flavor*). Single dots represent individual sessions for monkey M (mk M, green) and X (violet), with box and whisker plots indicating the median (central line), interquartile range (box), and standard deviation. (**C**) Predicted probability of choosing flavor J1 against the log ratio of probabilities associated with the two outcome flavors for each session collected in monkey M (left) and X (right). Dashed lines represent the average choice probabilities across sessions. Monkeys’ choice depended on the offered probability (sigmoid function) and the outcome flavor (different bias from session to session).

The two options were associated with different outcome flavors (juice 1 vs. 2, referred to as *different-flavor trials*) in half of the trials for monkey M and in 3 out of 4 trials for monkey X. In these trials, the probability was always different for the two outcomes in monkey M but could be either different or similar in monkey X (e.g., juice 1 at 70% vs. juice 2 at 70%). In the remaining trials (1/2 for monkey M and 1/4 for monkey X), the two options were associated with the same outcome flavor (e.g., juice 1 vs. juice 1, referred to as *same-flavor trials*) but with different probabilities. Same-flavor trials were not considered in the current analyses.

### Surgical procedures and neural recordings

Surgical procedures and details on neural recordings were previously described (Stoll and Rudebeck, 2023). In brief, monkeys were implanted with a titanium head restraint device and a form-fitted plastic recording chamber that contained a 157-channel semi-chronic microdrive system (Gray Matter Research, Bozeman, MT; **Fig. 2A**) housing glass-coated electrodes (1-2MΩ at 1kHz; Alpha Omega Engineering, Nazareth, Israel). The current dataset contained neurons recorded across a total of 72 and 86 independently moveable electrodes that targeted ventral frontal cortex (monkey M and X, respectively).

**Figure 2.**
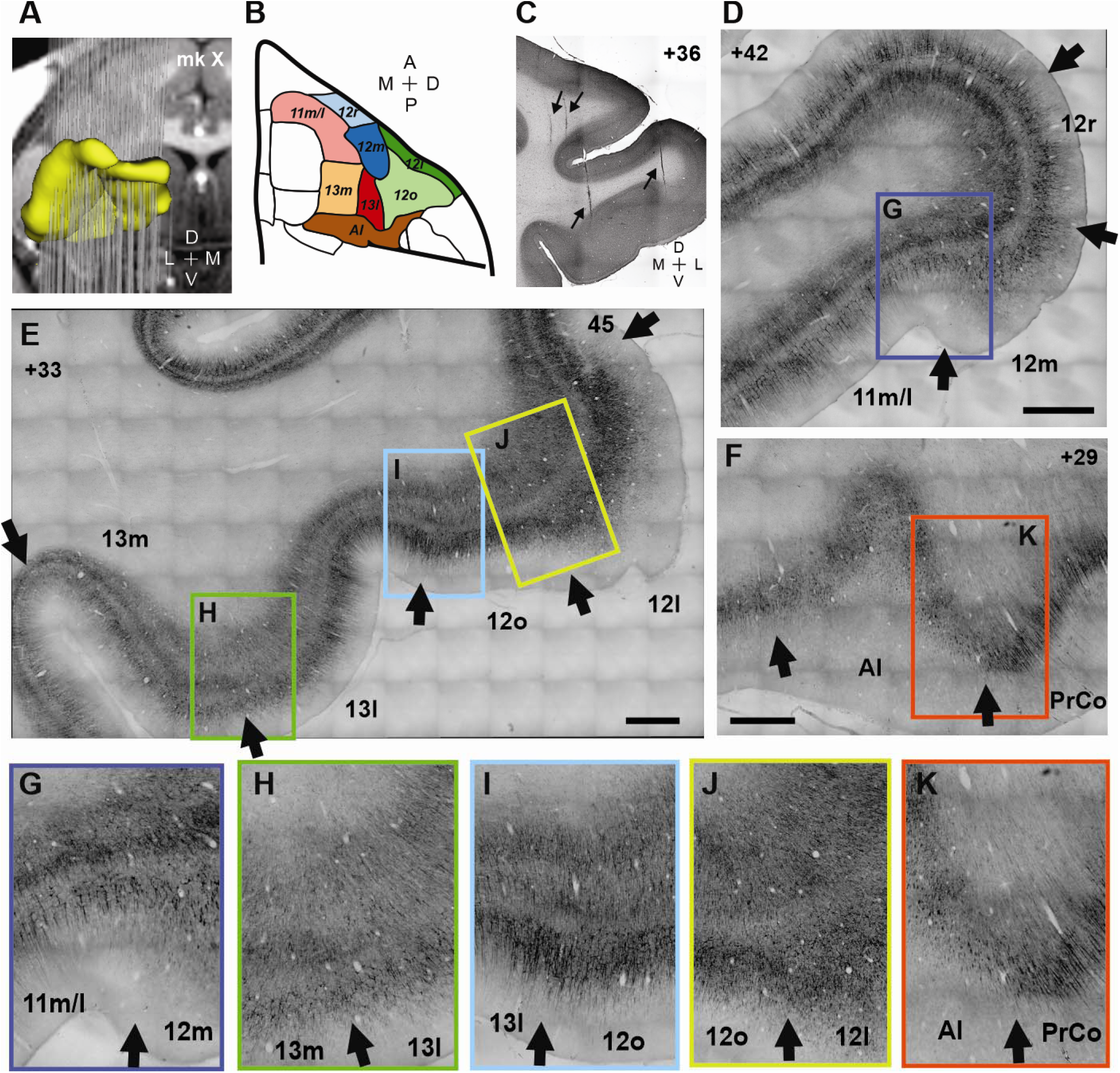
Recording locations and anatomical confirmation. (**A**) Model of the recording electrodes (grey) and extent of the targeted areas (yellow) aligned on the MRI on monkey X. (**B**) List of areas considered, represented on a ventral view of the macaque frontal cortex and colored as in following figures. (**C**) Coronal section of a Nissl staining at +36 interaural level in monkey M and highlighting 4 electrode tracks (small arrows). (**D-F**) SMI-32 stain at three antero-posterior levels (panel **D**: +42mm, **E**: +33mm and **F**: +29mm interaural) in monkey M. Large arrows represent transitions between subdivisions. Colored rectangles represent zoomed in portions displayed in following panels. (**G-K**) Detailed view of transitions between distinct subdivisions. Abbreviations: A=anterior, P=posterior, M/m=medial, L/l=lateral, D=dorsal, V=ventral, o=orbital, r=rostral, PrCo=precentral opercular cortex.

Recording locations were confirmed using several approaches. First, we recorded the cumulative depth of each electrode as they were being lowered and tracked the changes in background noise and electrophysiological activity suggestive of white/gray matter transitions. We also acquired CT images at different time points during the period where we were recording from ventral frontal cortex, which were then co-registered to post-operative MRIs to estimate the location of each electrode within the brain. Finally, after sacrifice and brain extraction, we captured and digitalized block-face images of every brain section which were later stained, as these showed clear marks of the electrodes’ track. Combined with microscope observations of histological stained sections (see below for more details) and using Free-D software (Andrey and Maurin, 2005), this allowed us to reconstruct the precise anatomical location of every electrode within the brain.

### Tissue preparation and immunohistochemistry

Following recordings, monkeys were deeply anesthetized and transcardially perfused with 4% formaldehyde in Phosphate-buffered saline. The brain was then extracted, post-fixed and cryo-preserved before being shipped to FD NeuroTechnologies, Inc (Columbia, MD) for tissue preparation and staining. Detailed methods were previously reported in Stoll and Rudebeck (2013). Briefly, the recorded brain hemispheres were further cryoprotected in solution before fast freezing in isopentane. Serial sections of 50μm were then cut coronally. Series of 4 consecutive sections were collected and stained separately (resolution of 200μm). Specifically, we stained every first section of each series using cresyl violet solution (Nissl staining), while the second and third series were processed for Calbindin (using mouse monoclonal anti-Calbindin-D-28K antibodies, 1:1000 dilution; Millipore Sigma, St. Louis, MO) and SMI-32 (using mouse Purified anti-Neurofilament H nonphosphorylated antibodies, 1:12000 dilution; Biolegend, San Diego, CA) immunohistochemistry.

### Defining neuroanatomical boundaries

Following the approach taken by Carmichael and Price (1994), we looked for variation in the composition of cortex across the ventral frontal cortex of each of our subjects on the Nissl, SMI-32, and calbindin-stained sections. We specifically chose these three stains as they qualitatively provided the greatest resolution to discern the areas of interest within ventral frontal cortex, namely parts of areas 11, 12, 13, and AI. Furthermore, recent anatomical analyses of the macaque frontal cortex by Rapan and colleagues have largely supported Carmichael and Price’s parcellation of ventral frontal cortex subdivisions (Rapan et al., 2023).

All sections were inspected at magnification between 2x and 20x, as required, using either Nikon or Zeiss light microscopes. The precise definitions of each area are provided in **Table 1** and examples of areal boundaries on stained sections are shown in **Fig. 2**. In each monkey we were reliably able to discern areas 13l, 13m, 12m, 12r, 12l, and 12o. While areas 11m and 11l could be reliably differentiated in one subject, they were harder to identify in the other. We therefore combined these two areas into one which we label 11m/l. We did not attempt to discern subdivisions of AI as the number of neurons in each would have been too few to meaningfully analyze.

**Table 1.** Structural characteristics used to define ventral frontal cortex subdivisions.

|  |  | Nissl | SMI-32 | Calbindin |
| --- | --- | --- | --- | --- |
| vIPFC | 12r | **Dysgranular**<br>Layer V not sublaminated<br>Vertical striations in III and V | Light stained cells in III and V | Light staining in superficial lamina (II and III) |
|  | 12m | **Granular**<br>Layer V laminated | Light stained cells in II and V | Dense stained cells in superficial lamina (II and III) |
|  | 12o | **Granular**<br>Weak layer IV and layer V not sublaminated | Light stained cells in II and V | Stained cells in both deep (V/VI) and superficial (II/III) lamina |
|  | 12l | **Granular**<br>Sharply laminated | Dense stained cells in III and V | Stained cells in both deep and superficial lamina |
| OFC | 11m/l | **Granular**<br>Aggregates of pyramidal cells in III and V | Dense stained cells in III | Dense staining and low vertical neurites in layer II/III |
|  | 13l | **Dysgranular**<br>layer V sublaminated | Two bands of dense stained cells in III and V | Many vertical processes<br>Dense stained cells in II/III |
|  | 13m | **Dysgranular**<br>no sublamination of III and V | Moderate, less intense staining of cells in III and V | Moderate number of stained vertical processes and cells |
| AI |  | **Agranular**<br>Layer V sublaminated | Dense stained cells in V | Dense stained cells in deep layers V and VI |

### Data processing

Data analyses were performed offline using custom Matlab scripts (Matlab version R2022a, MathWork Inc.) which are available at https://github.com/RudebeckLab/POTT-carto. Here, we focused our analyses on the instrumental task, specifically the response of neurons during trials where different outcome flavors were offered.

#### Behavior

Monkeys’ choices were analyzed using logistic regressions (function *fitglm* in Matlab), by fitting the odds of choosing one of the 2 flavors in trials where the two options were associated with different flavors using the following model:

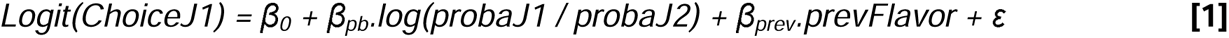

where *Logit(p) = log_e_(p/1 – p)*, the log ratio of probabilities associated with both outcome flavors, *probaJ1* and *probaJ2* (linear), and *prevFlavor* (J1 or J2, categorical) the last chosen outcome flavor in the preceding trial, β*_x_* the intercepts for each variable and ε the residuals. Note that the last chosen outcome flavor was independent of which task (single option, instrumental, dynamic) or condition (different- or same-flavor trials) was presented or whether the animal received it or not. In case no options were presented on the previous trial (break of fixation or no trial initiation), the previous chosen flavor was used. We extracted the significance and estimates for each variable across all sessions and computed the predicted choice probabilities (**Fig. 1C**) for a wider range of offer probabilities than were presented to the monkeys (from 10% to 90% by steps of 1%) with *prevFlavor* parameter set to the flavor J1 (function *predict* in Matlab).

#### Pre-processing of neurophysiological data

Neurons were included based on the quality of isolation only, with no response pattern or firing rate restrictions. Spiking activity for each trial was first smoothed using a 50ms Gaussian kernel before being averaged over 20ms bins. Neurons’ firing rates were aligned to multiple events across trials (central fixation, stimulus onset, response fixation, feedback and reward onsets). Our analyses were either performed on these sliding windows (e.g., single neuron ANOVAs) or applied on the average firing rate for each neuron around events of interest, notably the stimulus (100-700ms following stimulus onset) and reward (100-700ms following reward delivery) periods. Note that the reward period was redefined as being from 100 to 600ms when assessing whether information was maintained over time (see below and **Fig. 8**). This was done to keep time windows of maximal but similar length across events, with the feedback period being limited to 500ms before the reward was delivered. Given that monkeys rarely chose the lowest probability, and to ensure our neural encoding and decoding analyses are well-powered, we only considered 4 levels of chosen probabilities (30-50-70-90%). Results are provided for each monkey as well as combined and are indicated by individual symbols in the plots. When performing cross-condition decoding and subspace analyses (**Figs. 7** and **9**), we only considered neurons with an average firing rate greater than 1Hz (across the considered trials and time bins) and at least 5 trials for each considered condition.

#### ANOVA on single neuron responses

We assessed the tuning of neurons’ binned firing rates using two different ANOVAs. For this analysis, the firing rate for each neuron was min-max normalized across trials and time bins. First, we explained the variance in firing rate from each neuron and at every time bin using three-way ANOVAs which included the following factors: Chosen Flavor (2 levels, categorical), Chosen Probability (4 levels, 30-50-70-90%, linear) and Chosen Side (2 levels, categorical). We also assessed the neurons’ tuning to reward parameters using two-way nested ANOVAs which included: Reward delivery (2 levels, categorical) and Reward Flavor when delivered (3 levels, J1, J2 and none, categorical, nested under Reward delivery). Neurons were considered as encoding a task factor significantly if they discriminated that factor for 3 consecutive bins (covering a time period of 100ms) with a threshold of p<0.01. This led to a chance level of ∼5% of significant neurons for any given factor when considering the period before the stimulus onset (100-700ms following central fixation) (e.g., black line on **Fig. 3A**). We also extracted the omega-squared, a measure of the effect size, for each factor included in the ANOVAs. To assess differences between areas across our measures (proportion of significant neurons and effect size), we fitted generalized logistic (or linear) mixed-effect models (*fitglme* function in Matlab) with area (categorical) as fixed factor as well as monkey (2 levels, categorical) and session (categorical) as random factors (intercept only) (e.g., **Table 3-4**). We then applied multiple comparison correction using false discovery rate (FDR) at p<0.05.

**Figure 3.**
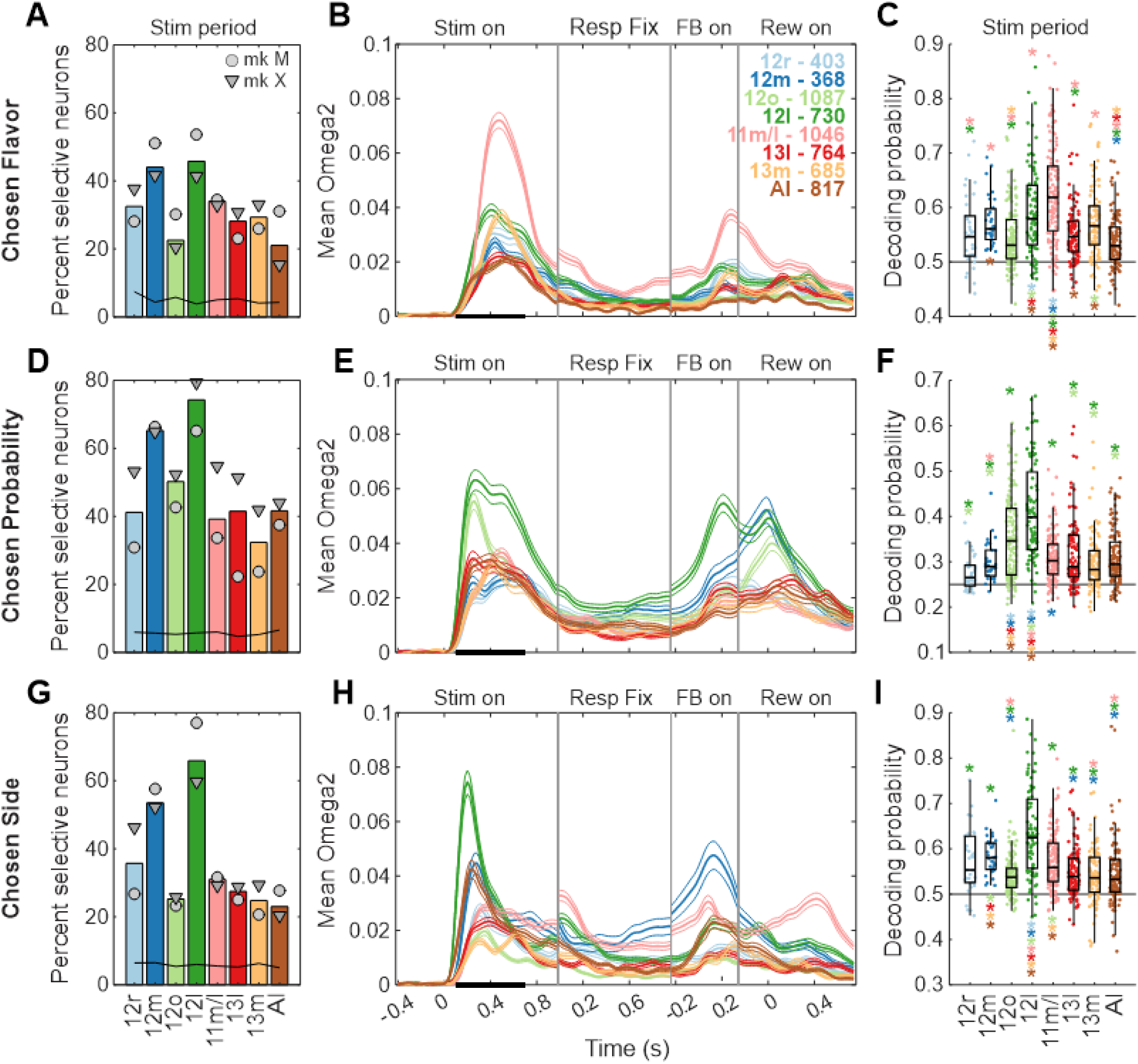
Encoding/Decoding of chosen flavor, chosen probability and chosen side during the stimulus period. (**A**) Percent of neurons across areas showing significant firing rate modulation with chosen outcome flavor (p<0.01 for 3 consecutive time bins) during the stimulus period (0.1 to 0.7s following stimulus onset, event ‘Stim on’). Black line shows the proportion of significantly modulated neurons during the pre-stimulus period (0.1 to 0.7s following central fixation) for each area. (**B**) Time-resolved average effect size for neurons with significant encoding of chosen flavor during the stimulus period (black bar), aligned to 4 successive events (Stimulus onset, Response fixation, Feedback onset and Reward onset). (**C**) Decoding performance for chosen flavor using simultaneously recorded neurons in each area. Each point represents a single session. Statistical significance was assessed using generalized mixed-effect models, with area as factor and monkey/session as random intercepts. Star’s locations (p<0.01 FDR corrected) indicate the direction of the effect (e.g., brown star – AI – under the yellow boxplot – 13m – means that AI decoding performance was significantly worse than 13m). The horizontal bar represents the theoretical chance level. (**D**-**F**) As in (A-C) but for chosen probability. (**G**-**I**) As in (A-C) but for chosen side. Circles and triangles represent data for monkey M and X respectively. See **Table 3** for statistical comparisons.

Next, we assessed the extent to which neurons signaling decision variables during the visual stimuli were reactivated in other periods of the task, including the feedback and reward periods (**Fig. 1**). To do this, we extracted the proportion of stimulus-related neurons that were also significantly tuned to the same parameter across three following 500ms time periods during the trial (Response: 100 to 600ms following response box fixation; Feedback: 0 to 500ms following feedback onset; Reward: 100 to 600ms following reward delivery). The beta coefficients associated with a given parameter across the stimulus period and each other time periods were correlated, and we extracted the across-neuron correlation coefficients as well as p-value (FDR-corrected at p<0.05).

#### Population decoding

We used population decoding methods to assess the strength of representations of the considered factors within both simultaneously recorded and pseudo-populations of neurons. For decoding on simultaneously recorded neurons, we only analyzed sessions with at least 4 neurons simultaneously recorded in the considered area. We first averaged the activity of all neurons from a single session across either the stimulus or reward periods before extracting a subset of similar number of trials for each category to avoid biasing classifiers (minimum of 5 trials for each category). This trial selection procedure and the following steps were performed 200 times. We then applied a principal component analysis (PCA) to extract the first 3 principal components. These were then used to decode the factor of interest by applying linear discriminant analysis (LDA) using 10-fold cross validation. The decoding performance on the testing sets were then averaged across the 10-fold cross-validations, before being averaged across the randomly sampled 200 trial permutations. Additionally, we assessed the significance of decoding performance by permuting the trial labels for each set of selected trials, resulting in 200 random permutations. Decoding was considered significant if less than 10 random permutation decoding performance were greater than the true average performance (one-tailed, p<0.05). As previously reported, we assessed differences between areas by fitting a generalized linear mixed-effect model (*fitglme* function in Matlab) with area as fixed factor as well as monkey and session as random factors (intercept only).

For decoding on pseudo-populations, we concatenated the activity of neurons recorded across sessions. To account for the different number of recorded neurons across areas, we ran the following analyses using multiple fixed numbers of neurons for each area (from 25 to 1000, see **Fig. 6**). For each run, we first applied a PCA on the time-averaged activity of the considered neurons, randomly sampled 200 times from our recording set, across random selection of trials (minimum of 10 trials for each category) and extracted the top 20 principal components, which were used to predict the parameter of interest using 10-fold cross-validated LDA. Decoding performance on the testing sets were then averaged across the cross-validations, before being averaged across the 200 trial/neuron-selection permutations. As before, we also permuted the trial labels 200 times to obtain random decoding performance, which was then statistically compared to the true decoding performance (one-tailed, p<0.05). We assess differences between areas using generalized linear mixed-effect model with area as fixed factor as well as monkey and the number of included neurons as random factors. Finally, we estimated the amount of information contained within neurons and populations by fitting a saturating function on the average decoding performance from the different ensemble sizes using:

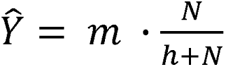

where Ŷ is the predicted decoding performance, *m* is the saturation level (referred to as the maximal information for a theoretical infinite population of neurons), *h* is the half-saturation point (characterizing the saturation rate and used to estimate the amount of information per neuron), and *N* the number of neurons included in the classifier.

#### Cross-condition pseudo-population decoding

We assessed the overlap in neural activity related to chosen flavor and chosen probability using a cross-condition decoding approach (e.g., (Wójcik et al., 2023), **Fig. 7B-C**). Compared to pseudo-population decoding, LDA classifiers were first trained to discriminate chosen probabilities on trials where a given flavor was chosen and then tested on trials where the other flavor was chosen (and vice versa). The same procedure was also used to train classifiers to decode the chosen outcome flavor on trials where a given probability was chosen and tested them on trials with a different chosen probability. We also ran the 10-fold cross-validated LDA classifiers without reducing dimensionality using PCA. As before, we performed cross-condition decoding using multiple fixed numbers of randomly selected neurons for each area (from 25 to 500), which was repeated 200 times (random neuron and trial selection). Cross-condition decoding performance was first averaged across the 2 classifiers (for chosen probability readout) or across all possible pairs of chosen probabilities (for chosen flavor readout), before being averaged across the 200 trial/neuron-selection permutations. We assessed significance using permutation testing, as previously described. We also compared the cross-condition performance with standard decoding performance obtained on the activity of the same considered neurons and trials but after permuting the non-decoded secondary parameter. As previously described, we also fitted saturating functions on the standard and cross-condition average decoding performance from the different ensemble sizes.

#### Principal component analyses (PCA) of neural reactivation

Here, the firing rate of each neuron was first z-scored and centered. For the qualitative comparisons shown in **Fig. 7A**, we extracted a random pseudo-population of 100 neurons for each area and applied a PCA on the average population activity across conditions (4 levels of chosen probability by the 2 chosen flavor, minimum of 5 trials per condition).

To assess whether chosen probability and reward representations shared neural spaces (**Fig. 9**), we implemented a PCA on pseudo-population activity from each subdivision of ventral frontal cortex. Specifically, to isolate the chosen probability subspace, we averaged the activity of each neuron across trials during the stimulus period (100-700ms following stimulus onset) for the 4 chosen probabilities. For the reward subspace, we averaged the activity of each neuron during the reward period (100-700ms following reward onset) in rewarded and non-rewarded trials. We then performed a PCA on each matrix, extracting in each case the eigenvectors of the first principal component. These eigenvectors were then used to project the average firing rate across time bins and each of the eight conditions (4 chosen probabilities by 2 reward conditions) onto the two identified neural subspaces. We repeated this procedure 100 times, each time randomly selecting a different pseudo-population of 100 neurons. We also ran a permutation for each of the 100 neurons selection procedure, in which we computed the average firing rate across the eight conditions after randomly permuting the trial labels, and before projecting the activity onto the previously isolated subspaces. For each time bin, we then compared the average Euclidean distance computed across all pairs of conditions (28 total comparisons across the 8 conditions) and for each of the 100-neuron selection sets with the 100 permutations (one tailed test). We considered differences to be significant when no permutations were greater than the truth (p<0.01) for 4 consecutive time bins. We further highlight only the significant time bins where these conditions were fulfilled in more than 50% of the neuron selection sets.

## RESULTS

### Task and behavioral performance

Two monkeys were trained to perform an instrumental choice task in which two options were presented simultaneously (**Fig. 1A**). Each option was composed of a central gauge, the level of which (more or less filled) signaled the probability that juice would be delivered if selected, and a colored frame indicating the potential juice flavor that would be delivered. Monkeys were free to select either option by fixating a response box located on each side of the screen. Following a feedback period in which both options were displayed again, a reward was delivered (or not) according to the monkeys’ choice (which defined the probability and flavor). We analyzed choices from 289 sessions (monkey M and X = 103 and 186 sessions) using a logistic regression model that included the log ratio of the probabilities associated with each outcome flavor. This model explained a large proportion of monkeys’ choice variability (**Fig. 1B**), revealing that the probability of receiving a reward had a strong influence on behavior. Outcome flavor also influenced monkeys’ choices, as highlighted by the variable shift in the sigmoid functions from one session to another (**Fig. 1C**, see also (Stoll and Rudebeck, 2023)).

### Recordings and anatomical confirmation

We recorded a total of 6,284 neurons across ventral frontal cortex. The trajectory of each electrode was determined based on post-mortem reconstructions of Nissl-stained coronal brain sections. These trajectories were then mapped to immunohistology stained sections and neurons were assigned to one of eight distinct ventral frontal cortex subdivisions based on the depth of each electrode at the time of recording (**Tables 1** and **2**). These subdivisions followed the detailed parcellations of this part of the frontal lobe reported by Price, Palomero-Gallagher, and colleagues (Carmichael and Price, 1994; Rapan et al., 2023).

**Table 2.** Number of recorded and analyzed neurons for each monkey. A session was included if 4 or more neurons from the same cytoarchitectonic area were simultaneously recorded.

| Area |  | Total number of Isolated Neurons |  | Number of neurons (Nb of Sessions) |  |
| --- | --- | --- | --- | --- | --- |
|  |  | mk M | mk X | *Figures 3-4* |  |
|  |  |  |  | mk M | mk X |
| vIPFC | 12r | 241 | 195 | 217 (27) | 186 (7) |
|  | 12m | 129 | 292 | 92 (4) | 276 (27) |
|  | 12o | 260 | 885 | 232 (27) | 855 (104) |
|  | 12l | 278 | 497 | 261 (27) | 469 (56) |
| OFC | 11m/l | 820 | 281 | 772 (95) | 274 (32) |
|  | 13l | 297 | 514 | 260 (35) | 504 (65) |
|  | 13m | 373 | 337 | 358 (51) | 327 (47) |
| AI |  | 347 | 538 | 296 (48) | 521 (62) |

### Representation of the decision variables during the stimulus period across ventral frontal cortex

Prior work has revealed the existence of a dissociation within ventral frontal cortex regarding the representation of outcome probability and outcome identity; the former relies on vlPFC whereas the latter relies on OFC (Rudebeck et al., 2017; Stoll and Rudebeck, 2023). Here we extend this dissociation by showing that neurons across cytoarchitectonic subdivisions within OFC and vlPFC do not represent information about outcome probability or identity uniformly (**Fig. 3**). This was evident across both encoding and decoding approaches at the level of single neurons (assessed using sliding window ANOVAs) and populations (assessed using LDA classifiers).

**Table 3.**
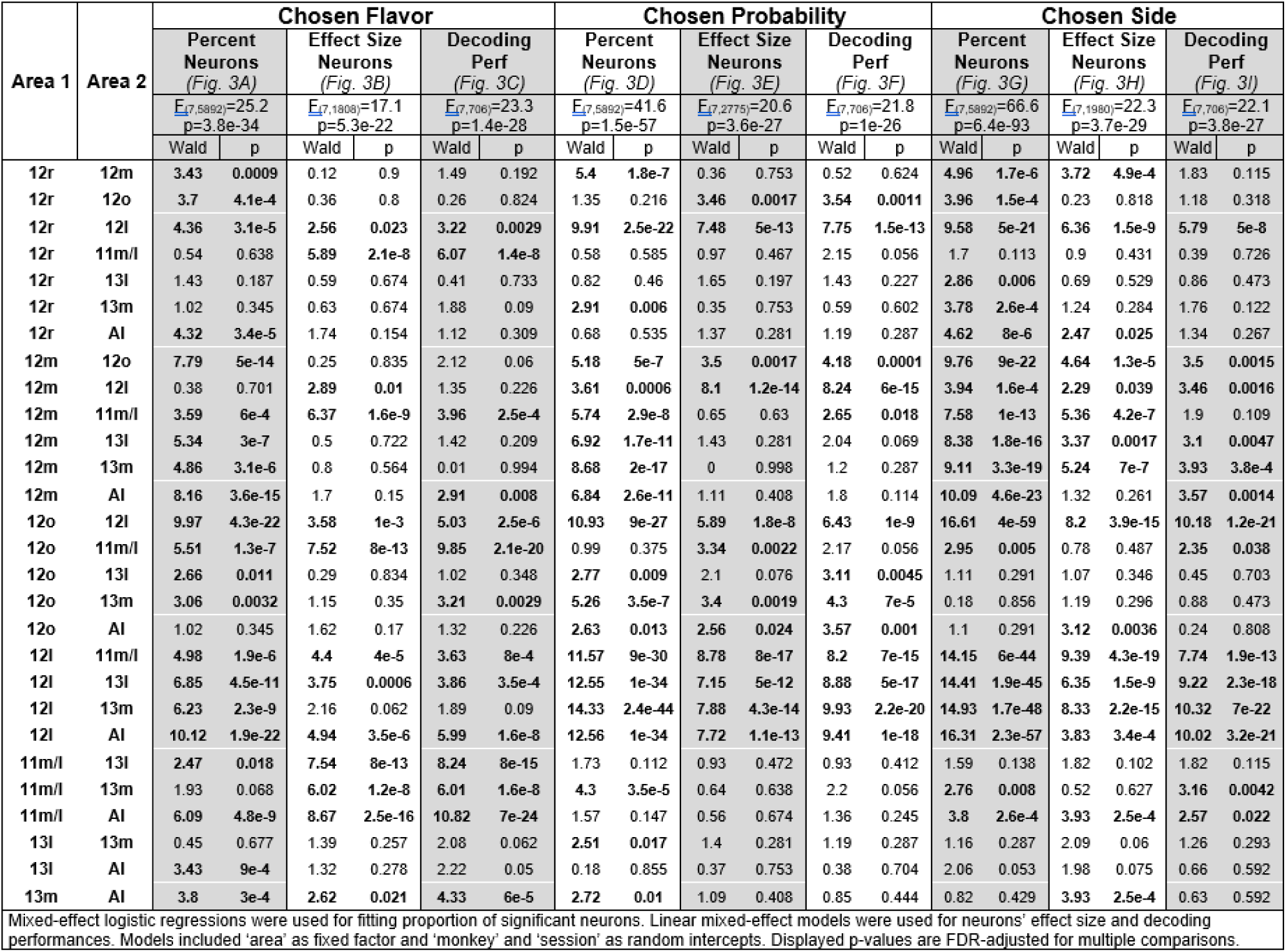
FDR-corrected multiple comparisons for area differences in percent of significant neurons, effect size and decoding performances for Chosen Flavor, Probability and Side.

During the stimulus period, 20 to 50% of neurons encoded the outcome flavor subjects would subsequently choose (**Fig. 3A**). 12m and 12l showed the highest proportion of neurons, followed by 12r and all OFC subdivisions, while 12o and AI showed the lowest proportion (mixed-effect logistic regression with FDR-correction for area comparisons in **Table 3**). Interestingly, however, a different pattern emerged regarding the effect size; a measure of how well the parameter of interest explained the variance in firing rate (**Fig. 3B**). Chosen flavor explained the activity of neurons in 11m/l, and to a lesser extent in 12l and 13m, better than in any other recorded areas. This was also true when assessing the ability to decode chosen flavor using the activity of simultaneously recorded neurons from single sessions. Here the highest decoding performance was observed within 11m/l compared to all other areas (**Fig. 3C**). Decoding performance was also higher in 12l compared to other parts of OFC and vlPFC, which exhibited relatively poor decoding performance.

A large proportion of neurons were also tuned to the chosen outcome probability following stimulus onset, with specific vlPFC subdivisions showing the highest proportion of neurons classified as encoding this decision variable compared to other areas (**Fig. 3D**). Neurons in both 12l and 12o that were classified as encoding chosen outcome probability also showed larger effect sizes than other areas, although the time course differed between these areas (**Fig. 3E**). Encoding of chosen probability was more transient in 12o while neurons in 12l maintained a strong representation until shortly before the decision was reported. The time course of the effect size in these neurons varied not only during the stimulus period but peaked again at later time points during the trial, notably during the feedback and reward periods. We will explore this apparent ‘reactivation’ of encoding in more detail in a later analysis. Finally, when we looked at how chosen probability could be decoded from simultaneously recorded neurons, we again found that the best decoding performance was found in 12l, followed by 12o (**Fig. 3F**). All other areas including 12r/m, all subdivisions within OFC, and AI showed relatively poor decoding performance.

We previously reported strong representations of the chosen response side in inferior frontal gyrus, and to a lesser extent in vlPFC (Stoll and Rudebeck, 2023). Here, we found that 12l showed the highest proportion of neurons (**Fig. 3G**), explained variance (**Fig. 3H**) and population decoding performance of chosen side (**Fig. 3I**) compared to other areas. It is worth noting that the representation of chosen side in 12l was more transient than the representation of other parameters. 12m also exhibited moderate representations of the chosen side. As others have reported previously (Padoa-Schioppa and Assad, 2006; Kennerley and Wallis, 2009), chosen side was weakly represented in OFC.

Altogether, we found that many neurons across ventral frontal cortex represented the chosen flavor and probability of the outcomes that would follow a specific stimulus, further supporting the involvement of this part of frontal cortex in the valuation stage of decision making (Noonan et al., 2017; Murray and Rudebeck, 2018; Stoll and Rudebeck, 2023). Representations were, however, not uniform across subdivisions and were dissociable based on encoding and decoding approaches. Of note, neurons in 12l exhibited the strongest and most diverse representation of the different parameters critical for the valuation of the options presented on each trial. By contrast, neurons in 11m/l more selectively coded for the chosen flavor whereas neurons in 12o represented the chosen probability.

### Representation of the decision variables during the reward period across ventral frontal cortex

In addition to the valuation of different potential choice options, ventral frontal cortex is also involved in assigning received rewards to chosen stimuli, a process known as credit assignment that is central to contingent learning (Walton et al., 2010; Chau et al., 2015; Noonan et al., 2017; Behrens et al., 2018). Notably, changes in neural activity in response to reward delivery have been repeatedly found in large swath of the brain, and the ventral frontal cortex is no exception (for example, (Thorpe et al., 1983; Tremblay and Schultz, 1999; Padoa-Schioppa and Assad, 2006; Kennerley and Wallis, 2009)). Thus, we next assessed whether there were any differences in how reward delivery influenced the activity of neurons within the 8 different subdivisions of ventral frontal cortex that we recorded from.

We found that the activity of a large proportion of neurons across ventral frontal cortex were modulated by reward delivery (**Fig. 4A**). All vlPFC subdivisions and AI showed a higher proportion of tuned neurons compared to OFC subdivisions (mixed-effect logistic regression with FDR-correction for area comparisons in **Table 4**). However, no clear differences were found between vlPFC and OFC subdivisions regarding effect size, with the exception of 13m, which exhibited the lowest explained variance of all areas that we recorded from (**Fig. 4B**). Population decoding performance was also generally high across almost all subdivisions, with 12o and 12r showing the highest performance while 13m showed the lowest (**Fig. 4C**). By comparison, the flavor of the reward that was received was represented in the activity of a smaller proportion of neurons (**Fig. 4D-E**), with only moderate differences in proportion of neurons or effect sizes between ventral frontal subdivisions. The population representation of the chosen flavor following reward delivery was stronger in 11m/l compared to 12r, 12o, 13l and AI, and marginally different in 11m/l compared to 12l and 12m (**Fig. 4F**). Interestingly, no differences were observed between 11m/l and 13m. Thus, while reward delivery and the delivered reward flavor were represented by a large proportion of neurons in ventral frontal cortex, there were differences in the degree of encoding. Notably, vlPFC areas showed the strongest representations of reward delivery, while area 11m/l and 13m exhibited the strongest representation of juice flavor.

**Figure 4.**
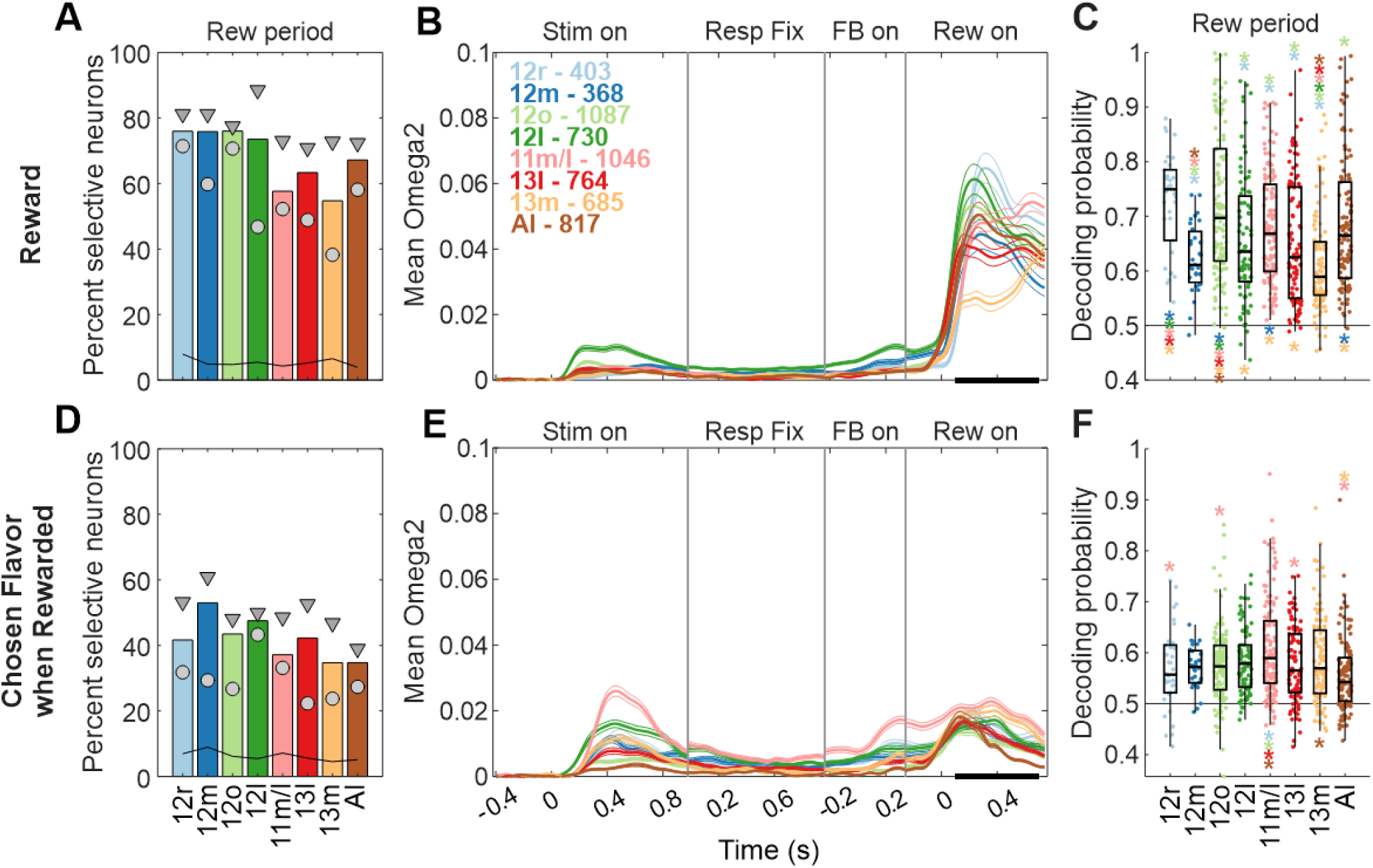
Encoding/Decoding of reward delivery and chosen outcome flavor during the reward period. (**A**) Percent of neurons across areas showing significant firing rate modulation whether reward was delivered or not during the reward period (0.1 to 0.7s following reward onset, event ‘Rew on’). Black line shows the proportion of significantly modulated neurons during the pre-stimulus period (0.1 to 0.7s following central fixation) for each area. (**B**) Time-resolved average effect size for neurons with significant encoding of reward delivery during the reward period (black bar), aligned to 4 successive events (Stimulus onset, Response fixation, Feedback onset and Reward onset). (**C**) Decoding performance for reward delivery using simultaneously recorded neurons in each area. Each point represents a single session. (**D**-**F**) As in (A-C) but for chosen flavor when rewarded. Conventions as in Fig. 3. See **Table 4** for statistical comparisons.

**Table 4.**
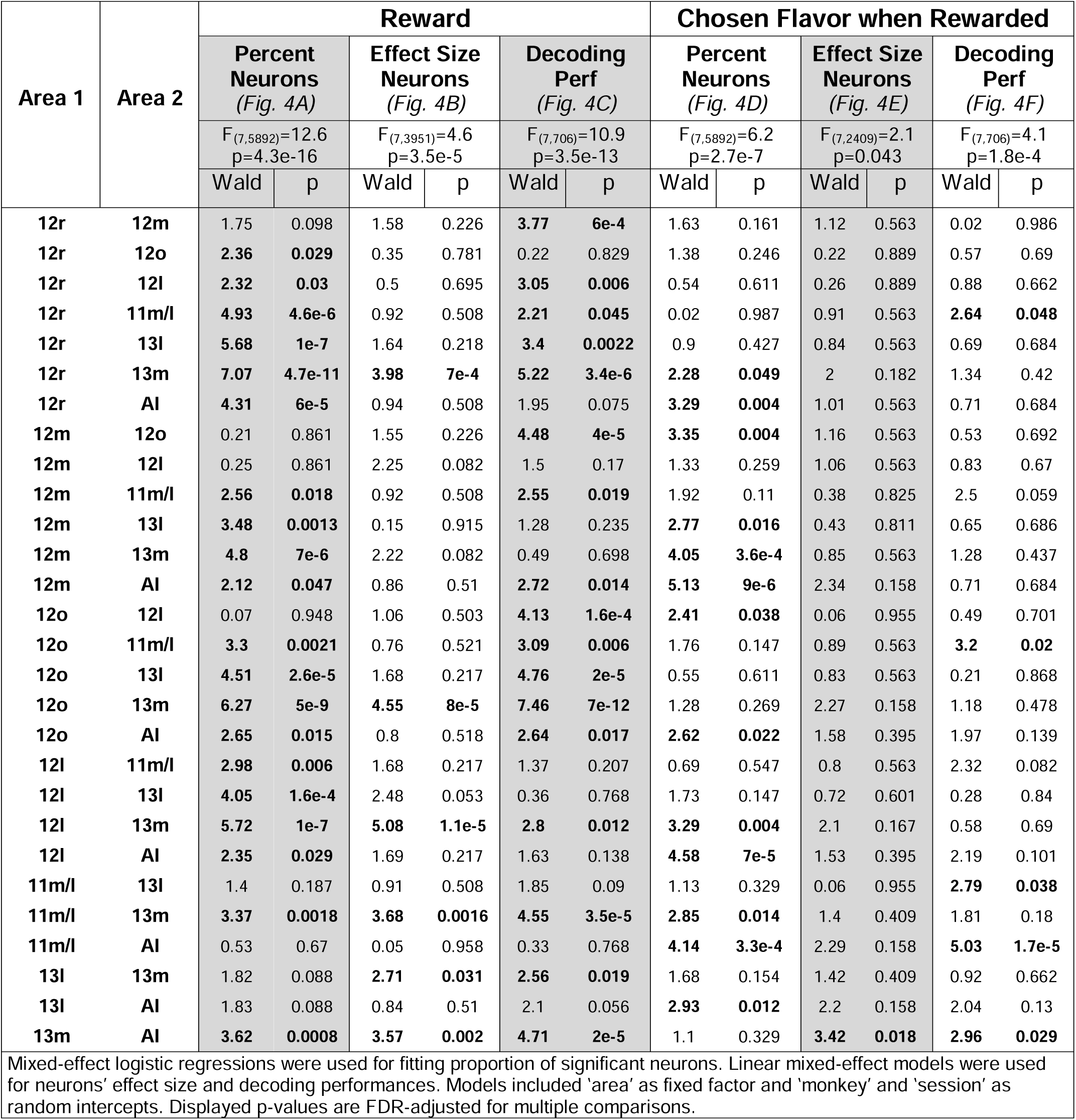
FDR-corrected multiple comparisons for area differences in percent of significant neurons, effect size and decoding performances for Reward and Chosen Flavor when Rewarded (related to. **Fig. 4).**

| Area 1 | Area 2 | Reward |  |  |  |  |  | Chosen Flavor when Rewarded |  |  |  |  |  |
| --- | --- | --- | --- | --- | --- | --- | --- | --- | --- | --- | --- | --- | --- |
|  |  | Percent Neurons<br>(Fig. 4A) |  | Effect Size Neurons<br>(Fig. 4B) |  | Decoding Perf<br>(Fig. 4C) |  | Percent Neurons<br>(Fig. 4D) |  | Effect Size Neurons<br>(Fig. 4E) |  | Decoding Perf<br>(Fig. 4F) |  |
| | | $F_{(7,5892)}=12.6$<br>$p=4.3e-16$ | | $F_{(7,3951)}=4.6$<br>$p=3.5e-5$ | | $F_{(7,706)}=10.9$<br>$p=3.5e-13$ | | $F_{(7,5892)}=6.2$<br>$p=2.7e-7$ | | $F_{(7,2409)}=2.1$<br>$p=0.043$ | | $F_{(7,706)}=4.1$<br>$p=1.8e-4$ | |
|  |  | Wald | p | Wald | p | Wald | p | Wald | p | Wald | p | Wald | p |
| 12r | 12m | 1.75 | 0.098 | 1.58 | 0.226 | **3.77** | **6e-4** | 1.63 | 0.161 | 1.12 | 0.563 | 0.02 | 0.986 |
| 12r | 12o | **2.36** | **0.029** | 0.35 | 0.781 | 0.22 | 0.829 | 1.38 | 0.246 | 0.22 | 0.889 | 0.57 | 0.69 |
| 12r | 12l | **2.32** | **0.03** | 0.5 | 0.695 | **3.05** | **0.006** | 0.54 | 0.611 | 0.26 | 0.889 | 0.88 | 0.662 |
| 12r | 11m/l | **4.93** | **4.6e-6** | 0.92 | 0.508 | **2.21** | **0.045** | 0.02 | 0.987 | 0.91 | 0.563 | **2.64** | **0.048** |
| 12r | 13l | **5.68** | **1e-7** | 1.64 | 0.218 | **3.4** | **0.0022** | 0.9 | 0.427 | 0.84 | 0.563 | 0.69 | 0.684 |
| 12r | 13m | **7.07** | **4.7e-11** | **3.98** | **7e-4** | **5.22** | **3.4e-6** | **2.28** | **0.049** | 2 | 0.182 | 1.34 | 0.42 |
| 12r | AI | **4.31** | **6e-5** | 0.94 | 0.508 | 1.95 | 0.075 | **3.29** | **0.004** | 1.01 | 0.563 | 0.71 | 0.684 |
| 12m | 12o | 0.21 | 0.861 | 1.55 | 0.226 | **4.48** | **4e-5** | **3.35** | **0.004** | 1.16 | 0.563 | 0.53 | 0.692 |
| 12m | 12l | 0.25 | 0.861 | 2.25 | 0.082 | 1.5 | 0.17 | 1.33 | 0.259 | 1.06 | 0.563 | 0.83 | 0.67 |
| 12m | 11m/l | **2.56** | **0.018** | 0.92 | 0.508 | **2.55** | **0.019** | 1.92 | 0.11 | 0.38 | 0.825 | 2.5 | 0.059 |
| 12m | 13l | **3.48** | **0.0013** | 0.15 | 0.915 | 1.28 | 0.235 | **2.77** | **0.016** | 0.43 | 0.811 | 0.65 | 0.686 |
| 12m | 13m | **4.8** | **7e-6** | 2.22 | 0.082 | 0.49 | 0.698 | **4.05** | **3.6e-4** | 0.85 | 0.563 | 1.28 | 0.437 |
| 12m | AI | **2.12** | **0.047** | 0.86 | 0.51 | **2.72** | **0.014** | **5.13** | **9e-6** | 2.34 | 0.158 | 0.71 | 0.684 |
| 12o | 12l | 0.07 | 0.948 | 1.06 | 0.503 | **4.13** | **1.6e-4** | **2.41** | **0.038** | 0.06 | 0.955 | 0.49 | 0.701 |
| 12o | 11m/l | **3.3** | **0.0021** | 0.76 | 0.521 | **3.09** | **0.006** | 1.76 | 0.147 | 0.89 | 0.563 | **3.2** | **0.02** |
| 12o | 13l | **4.51** | **2.6e-5** | 1.68 | 0.217 | **4.76** | **2e-5** | 0.55 | 0.611 | 0.83 | 0.563 | 0.21 | 0.868 |
| 12o | 13m | **6.27** | **5e-9** | **4.55** | **8e-5** | **7.46** | **7e-12** | 1.28 | 0.269 | 2.27 | 0.158 | 1.18 | 0.478 |
| 12o | AI | **2.65** | **0.015** | 0.8 | 0.518 | **2.64** | **0.017** | **2.62** | **0.022** | 1.58 | 0.395 | 1.97 | 0.139 |
| 12l | 11m/l | **2.98** | **0.006** | 1.68 | 0.217 | 1.37 | 0.207 | 0.69 | 0.547 | 0.8 | 0.563 | 2.32 | 0.082 |
| 12l | 13l | **4.05** | **1.6e-4** | 2.48 | 0.053 | 0.36 | 0.768 | 1.73 | 0.147 | 0.72 | 0.601 | 0.28 | 0.84 |
| 12l | 13m | **5.72** | **1e-7** | **5.08** | **1.1e-5** | **2.8** | **0.012** | **3.29** | **0.004** | 2.1 | 0.167 | 0.58 | 0.69 |
| 12l | AI | **2.35** | **0.029** | 1.69 | 0.217 | 1.63 | 0.138 | **4.58** | **7e-5** | 1.53 | 0.395 | 2.19 | 0.101 |
| 11m/l | 13l | 1.4 | 0.187 | 0.91 | 0.508 | 1.85 | 0.09 | 1.13 | 0.329 | 0.06 | 0.955 | **2.79** | **0.038** |
| 11m/l | 13m | **3.37** | **0.0018** | **3.68** | **0.0016** | **4.55** | **3.5e-5** | **2.85** | **0.014** | 1.4 | 0.409 | 1.81 | 0.18 |
| 11m/l | AI | 0.53 | 0.67 | 0.05 | 0.958 | 0.33 | 0.768 | **4.14** | **3.3e-4** | 2.29 | 0.158 | **5.03** | **1.7e-5** |
| 13l | 13m | 1.82 | 0.088 | **2.71** | **0.031** | **2.56** | **0.019** | 1.68 | 0.154 | 1.42 | 0.409 | 0.92 | 0.662 |
| 13l | AI | 1.83 | 0.088 | 0.84 | 0.51 | 2.1 | 0.056 | **2.93** | **0.012** | 2.2 | 0.158 | 2.04 | 0.13 |
| 13m | AI | **3.62** | **0.0008** | **3.57** | **0.002** | **4.71** | **2e-5** | 1.1 | 0.329 | **3.42** | **0.018** | **2.96** | **0.029** |

Mixed-effect logistic regressions were used for fitting proportion of significant neurons. Linear mixed-effect models were used for neurons' effect size and decoding performances. Models included 'area' as fixed factor and 'monkey' and 'session' as random intercepts. Displayed p-values are FDR-adjusted for multiple comparisons.

**Table 5.**
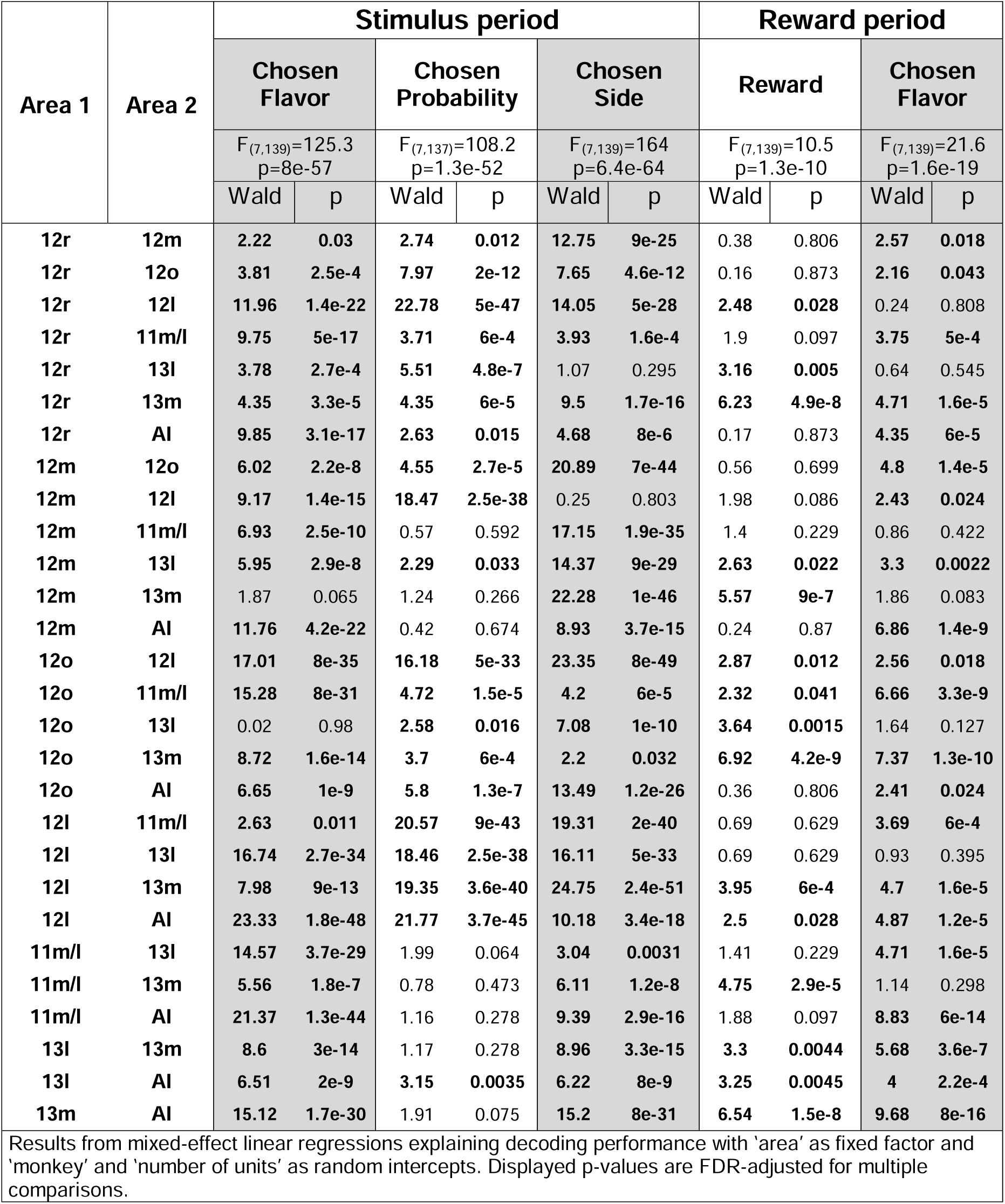
FDR-corrected multiple comparison for area differences in pseudo-population decoding performances for all parameters of interest (related to. **Fig. 6).**

| Area 1 | Area 2 | Stimulus period |  |  |  |  |  | Reward period |  |  |  |
| --- | --- | --- | --- | --- | --- | --- | --- | --- | --- | --- | --- |
|  |  | Chosen Flavor |  | Chosen Probability |  | Chosen Side |  | Reward |  | Chosen Flavor |  |
|  |  | F<sub>(7,139)</sub>=125.3<br>p=8e-57 |  | F<sub>(7,137)</sub>=108.2<br>p=1.3e-52 |  | F<sub>(7,139)</sub>=164<br>p=6.4e-64 |  | F<sub>(7,139)</sub>=10.5<br>p=1.3e-10 |  | F<sub>(7,139)</sub>=21.6<br>p=1.6e-19 |  |
|  |  | Wald | p | Wald | p | Wald | p | Wald | p | Wald | p |
| 12r | 12m | 2.22 | 0.03 | 2.74 | 0.012 | 12.75 | 9e-25 | 0.38 | 0.806 | 2.57 | 0.018 |
| 12r | 12o | 3.81 | 2.5e-4 | 7.97 | 2e-12 | 7.65 | 4.6e-12 | 0.16 | 0.873 | 2.16 | 0.043 |
| 12r | 12l | 11.96 | 1.4e-22 | 22.78 | 5e-47 | 14.05 | 5e-28 | 2.48 | 0.028 | 0.24 | 0.808 |
| 12r | 11m/l | 9.75 | 5e-17 | 3.71 | 6e-4 | 3.93 | 1.6e-4 | 1.9 | 0.097 | 3.75 | 5e-4 |
| 12r | 13l | 3.78 | 2.7e-4 | 5.51 | 4.8e-7 | 1.07 | 0.295 | 3.16 | 0.005 | 0.64 | 0.545 |
| 12r | 13m | 4.35 | 3.3e-5 | 4.35 | 6e-5 | 9.5 | 1.7e-16 | 6.23 | 4.9e-8 | 4.71 | 1.6e-5 |
| 12r | AI | 9.85 | 3.1e-17 | 2.63 | 0.015 | 4.68 | 8e-6 | 0.17 | 0.873 | 4.35 | 6e-5 |
| 12m | 12o | 6.02 | 2.2e-8 | 4.55 | 2.7e-5 | 20.89 | 7e-44 | 0.56 | 0.699 | 4.8 | 1.4e-5 |
| 12m | 12l | 9.17 | 1.4e-15 | 18.47 | 2.5e-38 | 0.25 | 0.803 | 1.98 | 0.086 | 2.43 | 0.024 |
| 12m | 11m/l | 6.93 | 2.5e-10 | 0.57 | 0.592 | 17.15 | 1.9e-35 | 1.4 | 0.229 | 0.86 | 0.422 |
| 12m | 13l | 5.95 | 2.9e-8 | 2.29 | 0.033 | 14.37 | 9e-29 | 2.63 | 0.022 | 3.3 | 0.0022 |
| 12m | 13m | 1.87 | 0.065 | 1.24 | 0.266 | 22.28 | 1e-46 | 5.57 | 9e-7 | 1.86 | 0.083 |
| 12m | AI | 11.76 | 4.2e-22 | 0.42 | 0.674 | 8.93 | 3.7e-15 | 0.24 | 0.87 | 6.86 | 1.4e-9 |
| 12o | 12l | 17.01 | 8e-35 | 16.18 | 5e-33 | 23.35 | 8e-49 | 2.87 | 0.012 | 2.56 | 0.018 |
| 12o | 11m/l | 15.28 | 8e-31 | 4.72 | 1.5e-5 | 4.2 | 6e-5 | 2.32 | 0.041 | 6.66 | 3.3e-9 |
| 12o | 13l | 0.02 | 0.98 | 2.58 | 0.016 | 7.08 | 1e-10 | 3.64 | 0.0015 | 1.64 | 0.127 |
| 12o | 13m | 8.72 | 1.6e-14 | 3.7 | 6e-4 | 2.2 | 0.032 | 6.92 | 4.2e-9 | 7.37 | 1.3e-10 |
| 12o | AI | 6.65 | 1e-9 | 5.8 | 1.3e-7 | 13.49 | 1.2e-26 | 0.36 | 0.806 | 2.41 | 0.024 |
| 12l | 11m/l | 2.63 | 0.011 | 20.57 | 9e-43 | 19.31 | 2e-40 | 0.69 | 0.629 | 3.69 | 6e-4 |
| 12l | 13l | 16.74 | 2.7e-34 | 18.46 | 2.5e-38 | 16.11 | 5e-33 | 0.69 | 0.629 | 0.93 | 0.395 |
| 12l | 13m | 7.98 | 9e-13 | 19.35 | 3.6e-40 | 24.75 | 2.4e-51 | 3.95 | 6e-4 | 4.7 | 1.6e-5 |
| 12l | AI | 23.33 | 1.8e-48 | 21.77 | 3.7e-45 | 10.18 | 3.4e-18 | 2.5 | 0.028 | 4.87 | 1.2e-5 |
| 11m/l | 13l | 14.57 | 3.7e-29 | 1.99 | 0.064 | 3.04 | 0.0031 | 1.41 | 0.229 | 4.71 | 1.6e-5 |
| 11m/l | 13m | 5.56 | 1.8e-7 | 0.78 | 0.473 | 6.11 | 1.2e-8 | 4.75 | 2.9e-5 | 1.14 | 0.298 |
| 11m/l | AI | 21.37 | 1.3e-44 | 1.16 | 0.278 | 9.39 | 2.9e-16 | 1.88 | 0.097 | 8.83 | 6e-14 |
| 13l | 13m | 8.6 | 3e-14 | 1.17 | 0.278 | 8.96 | 3.3e-15 | 3.3 | 0.0044 | 5.68 | 3.6e-7 |
| 13l | AI | 6.51 | 2e-9 | 3.15 | 0.0035 | 6.22 | 8e-9 | 3.25 | 0.0045 | 4 | 2.2e-4 |
| 13m | AI | 15.12 | 1.7e-30 | 1.91 | 0.075 | 15.2 | 8e-31 | 6.54 | 1.5e-8 | 9.68 | 8e-16 |

Results from mixed-effect linear regressions explaining decoding performance with 'area' as fixed factor and 'monkey' and 'number of units' as random intercepts. Displayed p-values are FDR-adjusted for multiple comparisons.

### Mixed selectivity within ventral frontal cortex

Numerous prior reports have shown that neurons across frontal cortex often exhibit “mixed selectively” in that they code for multiple task features (Rigotti et al., 2013; Fusi et al., 2016). We and others previously reported that vlPFC and OFC neurons are highly likely to exhibit mixed selectivity while maintaining orthogonal representations of the factors important for valuation (for instance, (Wallis and Miller, 2003; Stoll and Rudebeck, 2023). To establish if there were any differences in the degree of mixed selectivity between subdivisions of ventral frontal cortex, we conducted a similar set of analyses here.

Although mixed selectivity was observed in neurons across all subdivisions, 12m and 12l were unique in that most stimulus-related neurons recorded exhibited a high degree of mixed selectivity (**Fig. 5A**). Specifically, we found 472/730 (64.6%) of 12l neurons and 202/368 (54.9%) of 12m neurons exhibiting modulation of their firing rate by two or more parameters during the stimulus period (**Fig. 5B**). In fact, 12l showed the greatest proportion of neurons tuned to all three parameters that we considered in our analysis of neural activity (chosen flavor, probability, and side) compared to all other areas (FDR-corrected Chi-square tests, Chi2>12.4, p<9.1e-4), with 244/730 (33.4%) neurons. By comparison, all OFC subdivisions were characterized by less than 10% of such multimodal neurons (from n=108/1046 in 11m/l to n=41/685 in 13m). Thus, mixed selectivity was highest in parts of vlPFC, especially 12l.

**Figure 5.**
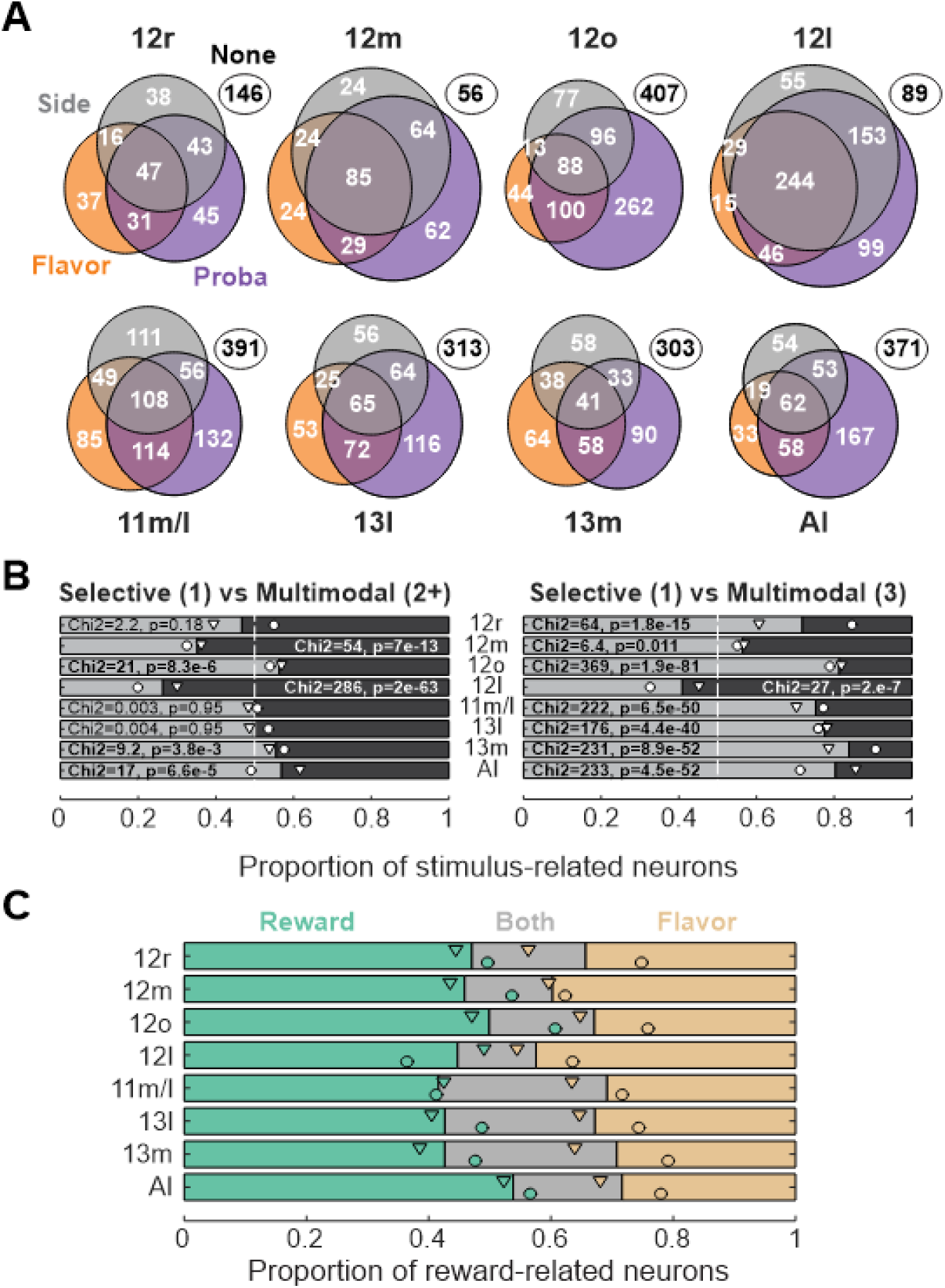
Mixed selectivity across ventral frontal cortex neurons. (**A**) Venn diagrams showing the numbers/proportions of neurons significantly representing chosen flavor (orange), chosen probability (purple), chosen side (grey) during the stimulus period. Number of non-selective neurons are displayed within the white ovals. (**B**) Proportion of neurons showing selective representation of a single parameter (light grey) compared to neurons showing multimodal integration (dark grey) across areas. Multimodal neurons are defined as representing 2 or 3 parameters (left panel) or all 3 parameters (right panel). Proportions were compared using Chi-square tests. Statistics and FDR-corrected p-values are displayed for each area (bold when significant). Circles and triangles represent the proportions for monkey M and X, respectively. (**C**) Proportion of neurons representing the reward (green), the chosen flavor in rewarded trials (yellow) or both (grey) during the reward period. Circles and triangles represent the proportions for monkey M and X, respectively.

At the time of the reward onset, a non-negligible proportion of neurons across areas represented both the reward receipt and its flavor (**Fig. 5C**; mixed-effect logistic regression, factor area, F_(7,4050)_=4.85, p=1.8e-5). In all of the OFC subdivisions, 24-28% of neurons exhibited this type of mixed selectivity. A different pattern emerged in parts of vlPFC, where the proportion of neurons exhibiting mixed selectivity for reward and flavor was lowest in 12l (72/563, 12.8%) compared to most other subdivisions (FDR-corrected post-hoc: 12l vs 12m/12r, W<2.1, p>0.11; 12l vs all other, W>2.95, p<0.013). It was also notable that area 12o and AI exhibited the highest degree of selective reward encoding neurons compared to OFC subdivisions (**Fig. 5C**, FDR-corrected Chi-square tests; AI vs OFC: Chi2>10.2, p<0.013; 12o vs OFC: Chi2>5.1, p<0.09). In summary, while 12l had the highest degree of mixed selectivity during the stimulus period, it showed the opposite pattern during the reward period.

### Representation of decision variables within pseudo-populations of ventral frontal neurons

In our previous analyses we were able to decode multiple task-relevant parameters using simultaneously recorded population of neurons across the different cortical subdivisions (**Figs. 3** and **4)**. Nevertheless, such an approach could be limited by the number of neurons we were able to simultaneously record from (median [min-max] population size of 5 [4-21] neurons) and might not reveal the full extent of ventral frontal cortex representations. We therefore assessed how well information could be decoded from the activity of neurons across ventral frontal cortex using pseudo-population of up to 1000 neurons, testing the robustness of our previous observations. Similar to what we observed before, clear differences in decoding performance were apparent across subdivisions (**Fig. 6**).

**Figure 6.**
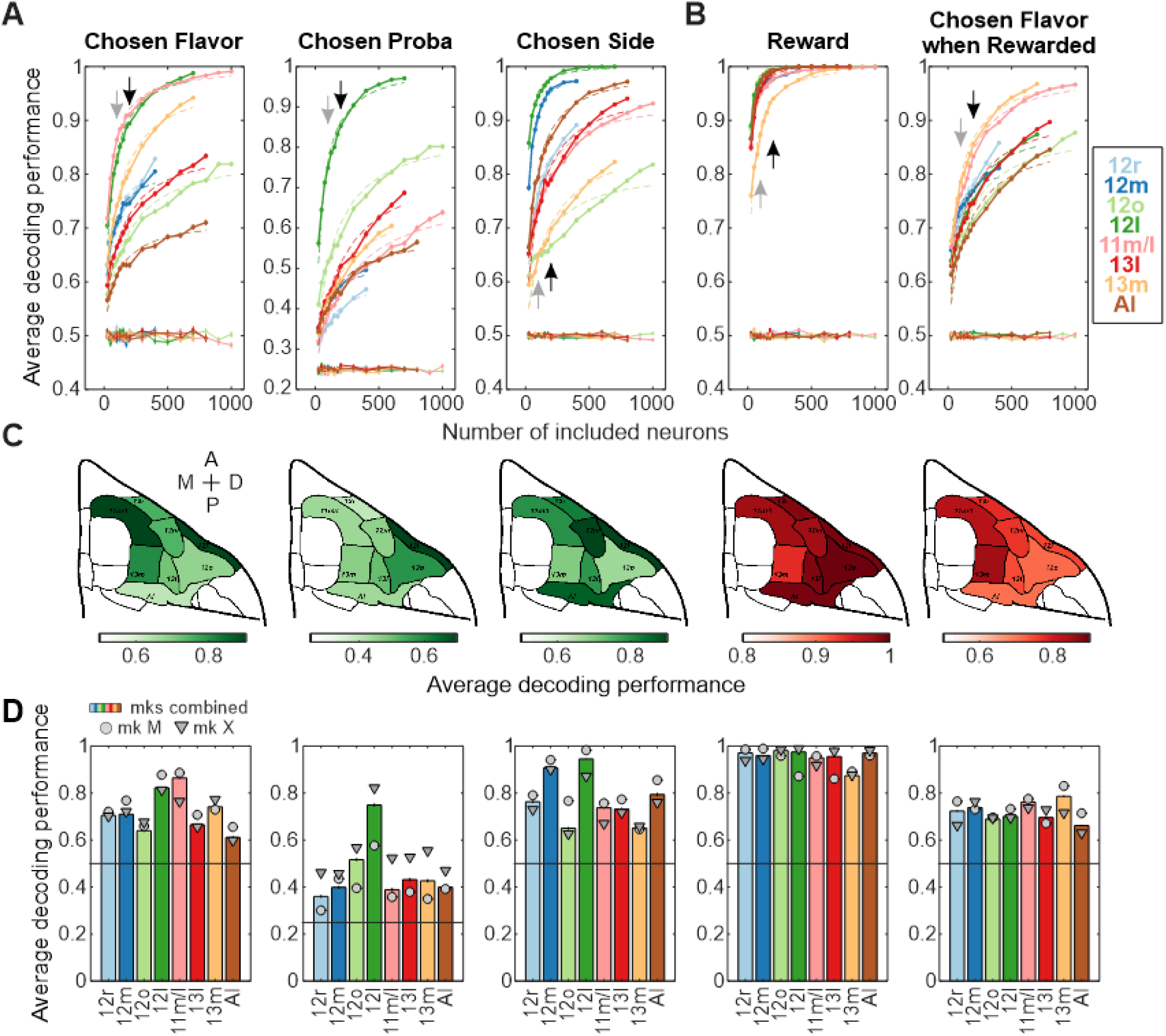
Pseudo-population decoding of tasks parameters across cytoarchitectonic areas. (**A**) Average decoding performance of the chosen flavor, chosen probability and chosen side (from left to right) during the stimulus period, using pseudo-populations of neurons from each isolated cytoarchitectonic area and against the size of the neuronal population considered. Dashed lines represent the fitted saturating functions on decoding performance while thin lines represent the average decoding performance across permutations. Black and grey arrows represent the numbers used for panel **C** and **D** respectively. (**B**) As panel A but showing the average decoding performance of reward receipt and chosen flavor when rewarded (left and right, respectively) during the reward period. (**C**) Average decoding performance using 200 neurons (black arrows in panel A-B) for the 5 considered parameters reported on a ventral view of the macaque frontal cortex (green and red colormap for the stimuli and reward periods respectively). (**D**) Average decoding performance using 100 neurons (grey arrows in panels A-B) for the 5 considered parameters, for each individual monkey (monkey M, circles; monkey X, triangles) as well as both combined (bars). Note that bars’ height did not necessarily fall in between the individual monkey points as the analyses were independent from each other. See **Table 5** for statistical comparisons.

Notably, we found that the population of area 12l neurons was unique compared to other areas in that it reached the highest level of decoding performance for all considered parameters during stimulus period (compare dark green lines to all others, **Fig. 6A, C-D**). After 12l, decoding performance was higher than other areas in 11m/l and 13m for the chosen flavor, while pseudo-populations of 12m and AI neurons more robustly represented the chosen side. Neurons in 12o had decoding performance that was higher than other areas for chosen probability, although at a level far lower than the performance of the 12l population (compare dark and light green lines for chosen probability, **Fig. 6A**). During the reward period, we found a strong representation of reward delivery in all areas, although the 13m population required more neurons to reach the highest level of performance compared to other areas (**Fig. 6B-D**). In contrast, pseudo-populations of neurons in area 13m as well as 11m/l had the highest decoding performance of the chosen flavor of the delivered reward. Contrary to what was found during the stimulus period, 12l pseudo-populations had the lowest decoding performance for outcome flavor. Such a distinction between stimulus and reward periods in area 12l potentially suggests a role for this area in the representation of value for choice, not the specific reward delivered, a point that we take up below.

Given the variation of representation across the different subdivisions of ventral frontal cortex, we next investigated how the variables that predominantly guided animals’ choices, chosen flavor and probability, were dynamically represented within the activity of pseudo-populations of neurons in each subdivision. Prior work has suggested that the diversity and dynamics of neural encoding are key components of the adaptive representation of task-relevant parameters and that this is required for flexible decision-making (Fusi et al., 2016). For this analysis we took the first 3 principal components from the neural activity of pseudo-populations of 100 randomly selected neurons from each subdivision and plotted the dynamic trajectories of activity associated with chosen probability and flavor 0 to 700ms after stimulus onset.

The different subdivisions of ventral frontal cortex exhibited qualitatively different neural population trajectories after the presentation of the visual stimuli (**Fig. 7A**). For example, the trajectory of activity from the top 3 principal components in 12o were clearly separated across the different levels of chosen probability (light to dark colors, **Fig. 7A**, top left) from shortly after the stimulus onset. By contrast, there was very little separation between the trajectories for chosen flavors (pink vs green). This indicates that dynamic representations in area 12o discriminate between the different levels of chosen probability, but not chosen flavor. 11m/l or 13m trajectories on the other hand displayed strong separation of chosen flavor with only a moderate separation of chosen probabilities (**Fig. 7A**, bottom row). Finally, 12l showed very distinct trajectories across both chosen flavor and probabilities such that all combinations of probability and flavor were clearly distinguishable (note the separation between different colors [flavor] and shading [probability], **Fig. 7A**, top right). Taken together, these observations echoed the differences in decoding performance between subdivisions that we previously found (**Figs. 3** and **6**) but also indicate that within each subdivision, chosen probability and flavor span distinct neural spaces.

**Figure 7.**
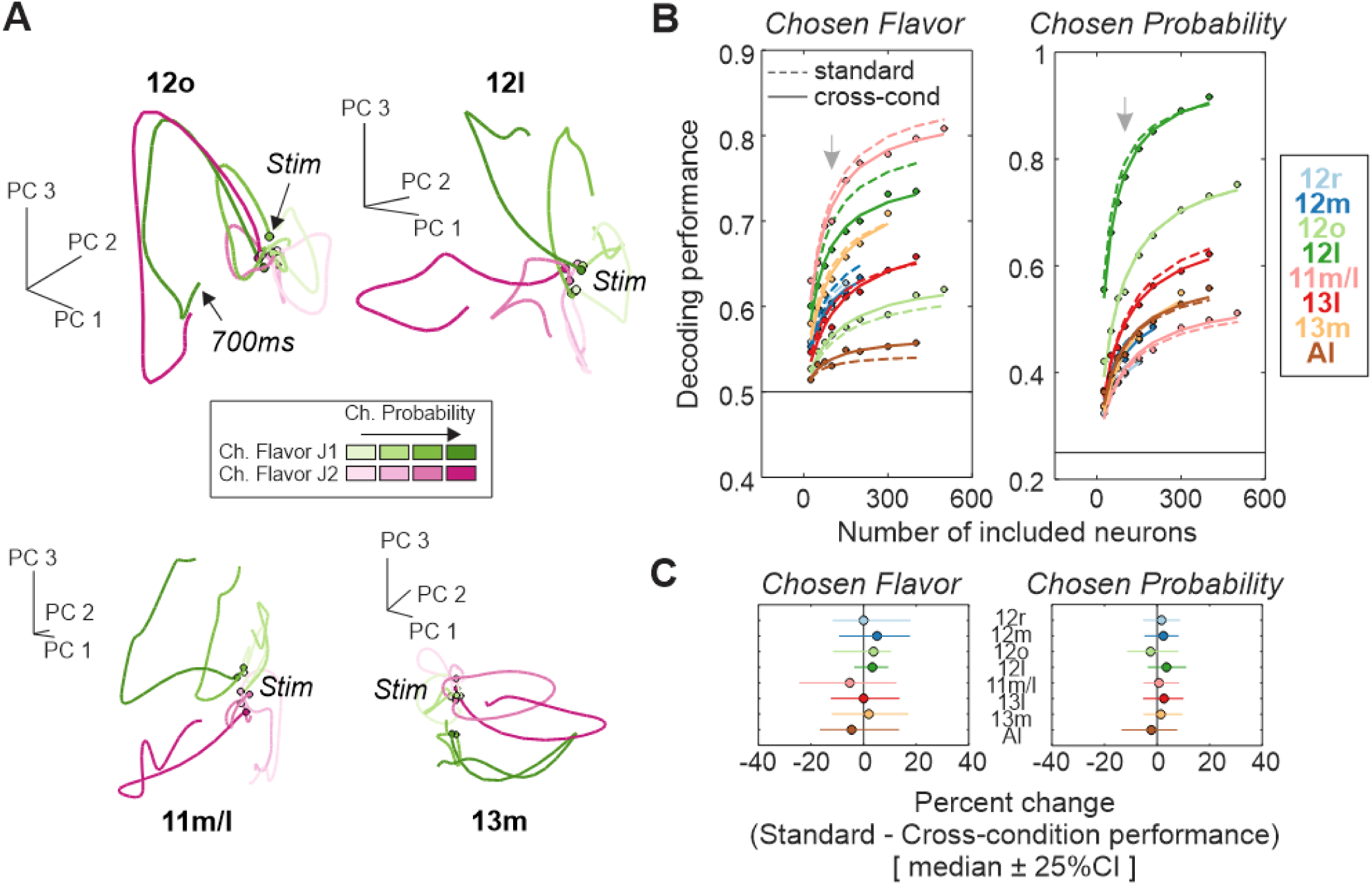
Orthogonal representations of chosen flavor and probability at the level of pseudo-population. (**A**) Top 3 principal components for pseudo-populations of 100 neurons recorded in 12o, 12l, 11m/l and 13m during the stimulus period. Green and pink curves represent the activity during chosen flavor J1 and J2 (respectively), while light to dark colors represent the increasing chosen probabilities (from 30 to 90%). Dots indicate the time of the stimulus onset. (**B**) Standard and cross-condition decoding performances of chosen f avor (left) and probability (right) using pseudo-populations of 25 to 500 neurons across all subdivisions. Dots represent raw cross-condition decoding performances while the lines represent the fitted saturated functions on either standard performances (dash) or cross-condition (plain). Grey arrows represent the number of included neurons used for panel C. (**C**) Median ± 25% CI percent change between standard and cross-conditions decoding performance (standard - cross-condition) when using 100 neurons across area and measures. Positive values indicate higher levels of performance for standard compared to cross-condition decoding.

To quantify the degree to which chosen probability and flavor were represented within distinct activity subspaces, we performed cross-condition decoding on the pseudo-populations of neurons from each subdivision (**Fig. 7B-C**). Here, we trained LDA classifiers to decode a given parameter (e.g., chosen probability) only using trials of a given class for another parameter (e.g., chosen flavor J1). Classifiers were then tested on the trials of the alternative class (e.g., chosen flavor J2) and we extracted the average decoding performance of our parameter of interest. If considering chosen probability decoding, high cross-condition decoding performance will only be observed if the weights associated with chosen probability representation for a given chosen flavor generalized from the alternative flavor, meaning chosen probability decoding is independent of the chosen flavor representation. Thus, high cross-condition could only be achieved if both chosen probability and chosen juice are represented within distinct neural subspaces (Bernardi et al., 2020; Wójcik et al., 2023).

We found that cross-condition decoding accuracies were often comparable to the performance of standard decoding across subdivisions for both primary task-relevant parameter, chosen probability or flavor (**Fig. 7B**). We found a greater decrease in performance for decoding chosen flavor using 12l pseudo-population compared to most other subdivisions. This difference between subdivisions is likely related to the fact that 12l is unique in that it exhibited the strongest representation of all three stimulus-related parameters, chosen flavor, probability, and side. Nevertheless, the average difference in performance between the two approaches was close to zero across all subdivisions (**Fig. 7C**, e.g. median change in chosen flavor decoding ranged from a 4.5% increase in performance for 11m/l to a 5.1% decrease in performance for 12m). Such an observation suggests that the representations of flavor and probability were orthogonal to each other.

### Dynamic encoding within ventral frontal cortex

As previously highlighted, neurons that were classified as encoding chosen flavor, outcome and side during the stimulus periods often exhibited similar levels of encoding at multiple times during each trial (**Fig. 3E**). For instance, neurons in areas 12o and 12l that encoded outcome probability during the stimulus period also had a peak of encoding during the reward periods (note the double peaks in dark and light green lines in **Fig. 3E**). Such dynamic representations might relate to the role of ventral frontal cortex in contingent learning as representations of stimulus-reward predictions are reactivated around reward delivery.

To explore this type of dynamic encoding, we first looked at whether single neurons in ventral frontal cortex encoded outcome flavor and/or probability at multiple points during each trial. The activity of the example neuron shown in **Fig. 8A** illustrates this type of dynamic encoding. The firing rate of this neuron was related to the chosen probability not only shortly after stimulus onset but also during the response, feedback, and reward parts of the trial. Mirroring the pattern of effects of this example neuron, we found that 20-70% of neurons encoding chosen probability or flavor during the stimulus period were ‘reactivated’ during other parts of the trial, most notably during the feedback and reward periods (**Fig. 8B**).

**Figure 8.**
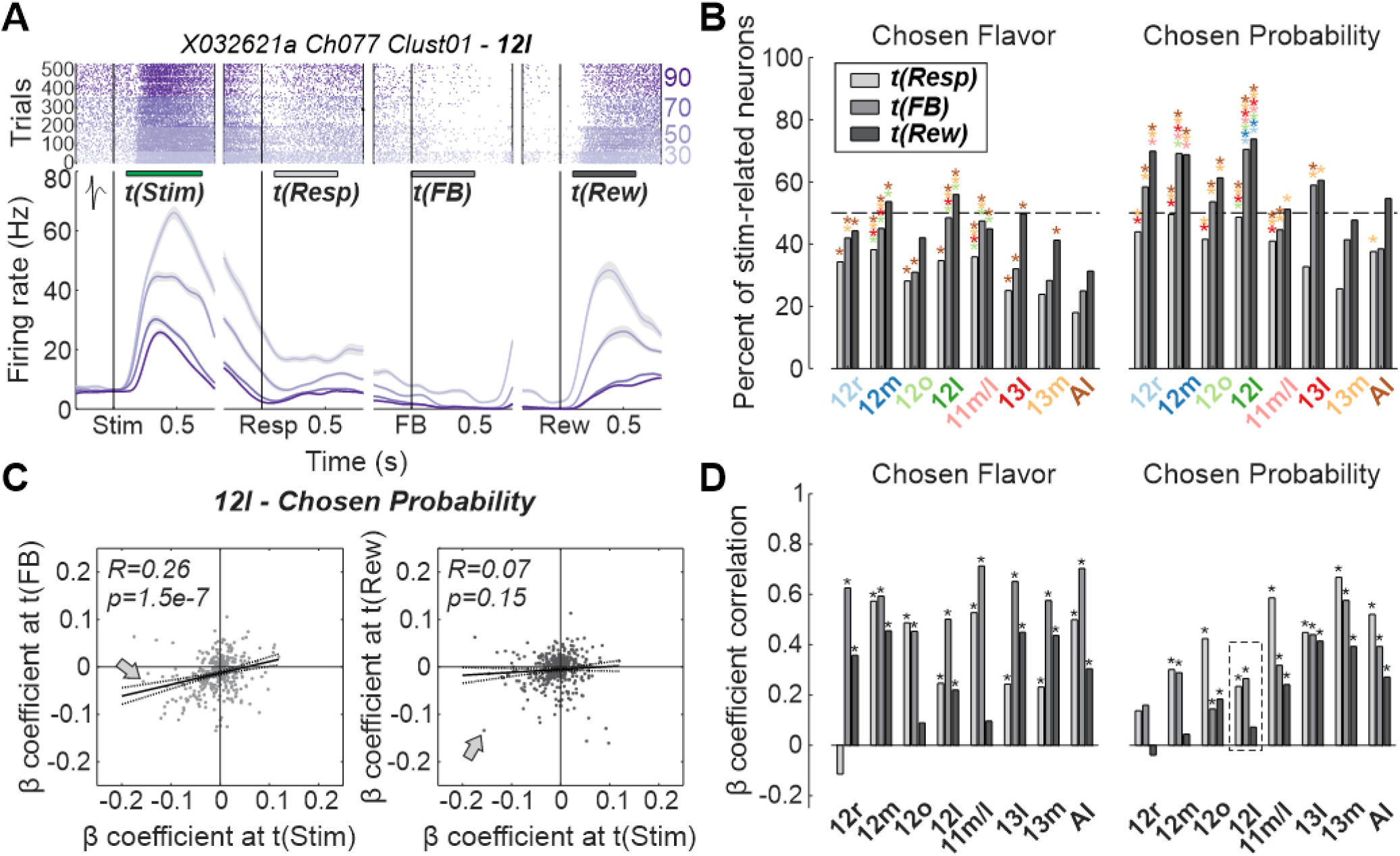
Temporally similar representations of chosen flavor and probability representations over time. (**A**) Raster plots and peri-stimulus time histograms of a 12l neuron aligned to multiple event onsets (stimulus, response, feedback, and reward) and for different chosen outcome probabilities (light to dark purple represent low to high probabilities respectively). Green/grey bars represented the time windows considered for the different events in the following panels. (**B**) Percent of significantly tuned neurons during the stimulus period which were also encoding the considered parameters (chosen flavor, probability, or side, from left to right) during the response fixation, feedback and reward periods (light to dark grey, respectively). Colored stars represent a significant difference in proportion (at FDR-corrected p<0.05) between two areas, the area where the star is located showing a greater proportion that the area indicated by the color of the star (e.g. the first brown star in the left panel on top of the proportion during response period for 12r means that this proportion is significantly greater than AI proportion). (**C**) Example correlations of the average beta coefficients related to chosen probability and obtained at different time window (left panel: stim vs FB; right panel: Stim vs Rew) across 12l neurons. Grey arrows highlight the coefficients associated with the neuron illustrated in panel A. (**D**) Correlation coefficients of beta coefficients for all areas and time window comparisons, as shown in panel B. Dashed square represent the correlations shown in panel C. Stars represent a significant correlation coefficient (at FDR-corrected p<0.05).

Regarding representations of chosen flavor, we found that a third of neurons maintained their encoding properties after monkeys made a response to select one of the options, while this proportion increased to 40-50% during the feedback and reward periods (**Fig. 8B**). Not all subdivisions exhibited the same rate of maintained chosen flavor encoding (mixed-effect logistic regressions, factor area, F_(7,2615)_>5.9, p<7.5e-7 across all time comparisons, **Fig. 8B**). Notably, there was a high proportion of neurons that represented flavor in both stimulus and reward periods within 12l and 12m (FDR-corrected post-hoc: 12l vs 13m/12o/AI, W>2.7, p<0.02; 12m vs 12o/AI, W>3.1, p<0.008). Within each area there were also notable differences in when the highest proportion of neurons showed similar encoding. For instance, there was a marked increase in the proportion of neurons in areas 13l and 13m that also encoded chosen juice in the reward period compared to the response or feedback periods. This indicates that there is a population of neurons in these areas which dynamically encode the chosen flavor over the course of the trial and that these neurons become active again around the time when the reward is likely to be delivered.

For chosen probability, a larger proportion of stimulus-related neurons continued to encode this decision variable later into the trial, reaching 50% across OFC and AI subdivisions and more than 70% within vlPFC subdivisions (mixed-effect logistic regressions, factor area, F_(7,3276)_>10.8, p<1.6e-13 across all time comparisons, **Fig. 8B**). Here we found that 12l showed the highest proportion of neurons encoding chosen probability at both the stimulus and feedback/reward periods compared to all other subdivisions (FDR-corrected post-hoc: 12l vs all areas, W>2.3, p<0.03 for both periods), closely followed by 12m neurons (stimulus-feedback periods; 12m vs 12r, W=1.9, p=0.07; 12m vs others, W>2.3, p<0.03).

The fact that the same neurons represent information across both stimulus and reward periods could be important for assessing whether the received reward meets expectations. It is nevertheless possible that neurons represent each task feature with either the same encoding scheme throughout these periods (as the example neuron in **Fig. 8A**) or using different encoding schemes. To assess this, we correlated the beta coefficients associated with a given task feature during the stimulus period and each of the other periods of the trial (examples for area 12l are shown in **Fig. 8C**). If a population of neurons is using the same encoding scheme across different parts of the trial, then there should be a positive correlation between the beta coefficients. If, however, different encoding schemes are being used then there should be a negative correlation or no correlation at all.

Overall, we found that most of the neurons represented chosen flavor in similar ways across periods, with significant positive correlations ranging from 0.4 to 0.7 (**Fig. 8D**, left panel). Of note, neurons in 11m/l and 12o, that represented chosen flavor at stimulus and reward periods to a high degree (note the high omega square for 11m/l and 12o in **Figs. 3-4**), showed high correlation between stimulus and response/feedback periods (R>0.45, p<3.9e-5), but not between stimulus and reward periods (R<0.1, p>0.22). In contrast to chosen flavor, correlations between beta coefficients from the stimulus and other periods of the trial were noticeably lower for chosen probability. We did, however, observe the same temporal correlation pattern in the two areas showing the best representations of chosen probability, namely 12l (**Fig. 8C**) and 12m (**Fig. 8D**, right panel) (stimulus vs response/feedback: R>0.23, p<1.2e-4; stimulus vs reward: R<0.07, p>0.15). In summary, the differences in representations of task-related parameters over time indicates that the encoding properties of neurons within ventral frontal cortex is dynamic. Indeed, we found that neurons maintained their tuning properties to different aspects of expected reward but did so with distinct activity patterns over time. Such a pattern of ‘reactivation’ of encoding potentially indicates that a given neuron could be involved in the assessment and updating of the choice options.

### Shared neural subspaces during stimulus and reward periods

At the level of single neurons, large proportions of vlPFC neurons represented chosen probability during both stimulus and reward periods, although with different encoding scheme over time. It remains an open question whether population activity subspaces identified during the stimulus period for chosen probability relate to the subspaces that were apparent during the reward period. One possibility is that representations of reward/no reward correspond to a binary transformation of the stimulus-related chosen probability representations, which would both exist within a similar neural space. Such a subspace might be important for updating specific reward-probability representations. Alternatively, chosen probability and reward representations could co-exist within distinct and separable neural activity subspaces, facilitating the readout of both types of information by other areas. To assess the temporal evolution of these representations and test our two hypotheses, we projected the pseudo-population activity over time onto the first component of 1) a chosen probability subspace extracted during the stimulus period and 2) a reward subspace extracted during the reward period (**Fig. 9A**). We then computed the time-resolved average Euclidean distance between every pair of conditions (4 chosen probabilities by 2 reward flavor condition) depending on which subspace the activity was projected onto (**Fig. 9B**). Here, larger distances represent a greater ability to discriminate probabilities and/or reward flavors.

**Figure 9.**
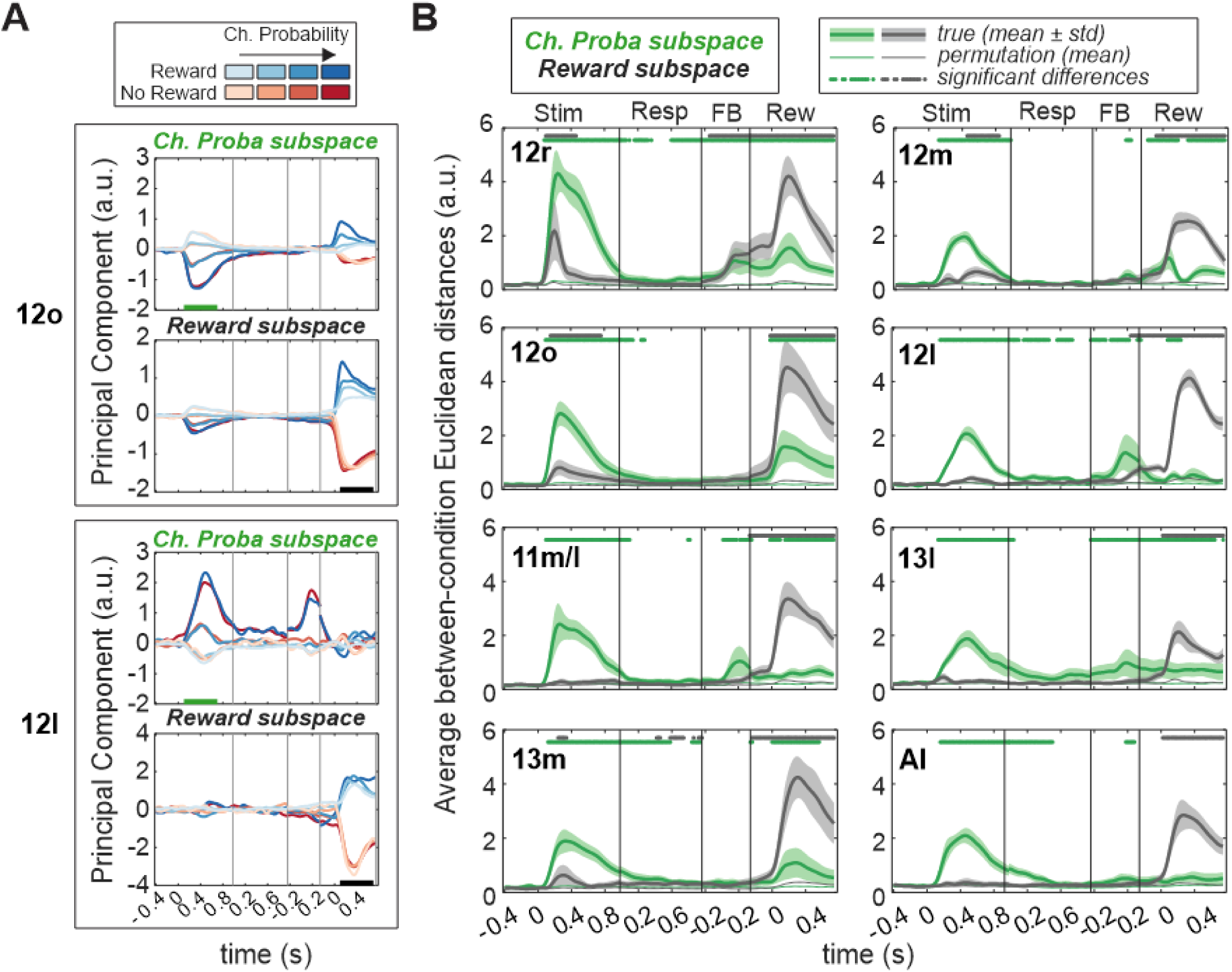
Temporal evolution of chosen probability and reward information across pseudo-populations. (**A**) Example of trial-averaged pseudo-population activity in 12o (top) and 12l (bottom) over time when projected onto the stimulus-related probability subspace (green) or the reward subspace (grey). Chosen probability levels are coded using light to dark colors while reward/no reward conditions are in blue/red respectively. Each pseudo-population was composed of a random selection of 100 neurons. Green and black bars on x-axis represent the time window used to derive the chosen probability and reward subspaces respectively. (**B**) Average Euclidean distance between all chosen probabilities and reward conditions across time and depending on whether the pseudo-population activity was projected onto the chosen probability (green) or reward (grey) subspace. Thick lines represent the average of the 100 neuron-selection permutations while shadings represent the standard deviation. Thin lines represent the average Euclidean distance when trial labels were permuted. Dots represent significant differences between the true distance and permutations, using a threshold at p<0.01 in more than 50% of the random neuron-selection sets.

In the pseudo-population of 12o neurons, probability and flavor information were contained within overlapping subspaces as evident by the two peaks of discrimination at the time of the stimulus and reward across each subspace (**Fig. 9A-B**). While this pattern was also seen in 12r, 12m and 13m, it was absent in 12l, 13l, and AI (note the lack of two peaks for stimulus and reward periods). In these latter subdivisions, the chosen probability subspace was prominent during the stimulus (gray peak in stimulus period) while the information content shifted toward the reward subspace during the reward (green peak in reward period). Such a pattern implies that the chosen probability and reward representations were largely separate or ‘disentangled’ in 12l and AI. Overall, the population representations within ventral frontal cortex subdivisions evolved differently over time and appear to indicate that different subdivisions are supporting different computations.

## DISCUSSION

Combining high-density single neuron recordings with precise anatomical reconstructions, we report that there is significant variation within ventral frontal cortex neural representations. Specifically, neurons in OFC area 11m/l were highly tuned to the outcome flavor at the time of the choice (**Figs. 3** and **6**). Neurons in more posterior OFC subdivisions 13m or 13l were more weakly tuned to outcome flavor although area 13m had the strongest representations of flavor at the time of reward delivery (**Figs. 4** and **6**). By contrast, neurons in vlPFC area 12l not only encoded outcome flavor, but also exhibited strong representations of outcome probability and choice direction (**Figs. 3** and **6**). In addition, neurons in area 12l also exhibited a high degree of mixed selectivity when stimuli were presented (**Fig. 5**) and this encoding was associated with distinct neural subspaces at the population level (**Figs. 7** and **9**). This potentially suggests a role for 12l in the integration of multiple decision variables during valuation to guide choice. This contrasted with vlPFC area 12o which showed selective representations of outcome probability (**Figs. 3** and **5**) as well as reward (**Fig 4**). Notably, stimulus- and reward-based representations existed within a shared neural subspace in 12o (**Fig. 9**), potentially indicating a neural mechanism for the putative role of this area in associating stimuli with rewards during learning. Taken together, our findings provide evidence that representations within ventral frontal cortex are not spatially or functionally uniform and indicates that each subdivision might contribute to specific cognitive processes.

### OFC and the coding of the specific outcome qualities

We previously reported that OFC neurons exhibited stronger representations of outcome flavor compared to vlPFC neurons (Stoll and Rudebeck, 2023). However, neurons across OFC subdivisions did not represent information uniformly. We observed strong chosen flavor representation in 11m/l neurons during both stimulus and reward periods, while 13m more strongly represented the chosen flavor when subjects received a reward. OFC area 13l and AI, on the other hand, exhibited low levels of tuning across both time periods.

Prior work has highlighted a role for OFC in valuation processes, especially when subjects are making value-based decisions (Thorpe et al., 1983; Tremblay and Schultz, 1999; Padoa-Schioppa and Assad, 2006; Kennerley and Wallis, 2009; Kennerley et al., 2009; Bouret and Richmond, 2010; Kobayashi et al., 2010; Rudebeck et al., 2013a; Pastor-Bernier et al., 2019; Stoll and Rudebeck, 2023). Regarding reward properties more specifically, neurons in both OFC areas 11 and 13 have been reported to respond to reward-predicting stimuli, both during reward expectation and after receipt, often discriminating between different outcome flavors in relation with animals’ preferences (Thorpe et al., 1983; Tremblay and Schultz, 1999; Padoa-Schioppa and Assad, 2006). However, this seminal work did not specifically look for whether distinct subdivisions of OFC represented value or flavor information differently. This was either because recordings focused on a specific subdivision (for instance area 13m in Padoa-Schioppa and Assad, 2006) or because the cytoarchitectonic area where recordings were made was not characterized (for instance, Thorpe et al., 1983).

Although not related to reward flavor, a small body of work supports the existence of antero-posterior gradient in OFC regarding the degree of information representations related to reward features (Sescousse et al., 2010; Klein-Flügge et al., 2013; Rich and Wallis, 2017). For example, Rich and Wallis reported stronger encoding of the expected reward size in high-gamma activity in more anterior compared to posterior parts of OFC (Rich and Wallis, 2017). This is somewhat consistent with our observation that OFC area 11m/l better represented the reward flavor during the stimulus period compared to areas 13m and 13l. The difference in encoding that we found also potentially concords with an inactivation study that highlighted a functional dissociation between areas 11 and 13 (Murray et al., 2015). Specifically, inactivation of area 13, but not area 11, impaired the updating of the sensory-specific values associated with a stimulus. By contrast, inactivation of area 11, but not area 13, impaired the ability of macaques to use the current value of a specific reward to appropriately guide goal selection. In the present task, subjects had previously learned the specific stimulus-flavor associations and used juice flavor to guide their choices (**Fig. 1**). Thus, stronger encoding of the chosen stimulus in area 11m/l compared to area 13 would fit with this role for area 11 in goal selection. This specific role in utilizing, as opposed to updating, reward flavor is further reinforced by the findings that representations of juice flavor in area 11m/l during the stimulus and reward periods exhibited low correlations (**Fig. 8**) and occupied largely separable neural subspaces (**Fig. 9**).

In contrast to area 11m/l, neurons in 13m were more specifically tuned to the flavor of the chosen outcome at the time of the reward delivery. Such a pattern of encoding in this area could represent a mechanism for the updating of specific stimulus-values based on the flavor of the received reward (Murray et al., 2015). Further, if area 13m is involved in updating stimulus-reward associations, then it could be expected that neural activity in this area at the time of reward should be more related to representations at the time of stimulus presentation. This is exactly what we found. In area 13m, but not 11/ml, the subspaces of activity activated at stimulus presentation overlapped with those during the reward period (**Fig. 9**). In addition, there was a high degree of similarity between the encoding of juice flavor in stimulus and reward periods in this area (**Fig. 8**). Further determining how these representations differ between parts of OFC, and what encoding in area 13l and AI are most related to, are questions that we were not able to fully address here, potentially as our task design was not diverse enough. Irrespective of this, our findings add to the growing body of work in humans and animals highlighting the diversity of representations in OFC related to guiding choices and updating representations (Padoa-Schioppa and Assad, 2006; Kennerley and Wallis, 2009; Klein-Flügge et al., 2013; Stalnaker et al., 2014; Rich and Wallis, 2016; Howard and Kahnt, 2017).

### Dissociable representations within vlPFC subdivisions

Apart from neuroimaging studies that have identified distinct face processing area in macaque vlPFC (Tsao et al., 2008), few studies have highlighted differences in encoding within this part of prefrontal cortex. We found that neurons in 12l were highly likely to encode task-relevant parameters during the stimulus period (**Fig. 3**), often representing more than a single parameter (**Fig. 5**). Such encoding is maybe not surprising given that area 12 receives projections from visual areas in inferior temporal cortex and a higher density of somatosensory inputs compared to other ventral frontal cortex subdivisions (Barbas, 1988). Despite this, we found that the representation of decision variables existed within distinct neural subspaces (**Figs. 7** and **9**). Compared to other vlPFC subregions, 12l is more strongly connected, anatomically and functionally, to lateral prefrontal cortex, notably area 45 and 46v (Saleem et al., 2014; Rapan et al., 2023). These more dorsal parts of lateral frontal cortex are more closely associated with guiding attention and actions towards the goal of a decision and are directly connected to premotor areas (Miller and Cohen, 2001; Kennerley et al., 2009; Cai and Padoa-Schioppa, 2014). Thus, such separate but high dimensional representations in 12l could be used by downstream areas to bias motor behavior and promote the selection of a specific action/motor plan.

Compared to neighboring subdivisions of area 12, representations in 12o were characterized by highly selective stimulus-related probability representations (**Figs. 3** and **5**) and coding of reward delivery (**Fig. 4** and **6**). Notably, stimulus probability and reward delivery subspaces overlapped in this area (**Fig. 9**). Such a pattern potentially indicates that, of the areas within ventral frontal cortex, 12o may be the most specialized for reward-based learning. Our findings thus provide single neuron and mechanistic insight into prior neuroimaging (Zald et al., 2005; Chau et al., 2015; Jocham et al., 2016) and interference approaches (Noonan et al., 2017; Rudebeck et al., 2017; Folloni et al., 2021) in humans and macaques that has emphasized a role for area 12o in contingent learning. While the present task does not specifically require stimulus-outcome associations to be updated, local fluctuations in reward history as well as changes in preference for the juice flavors (Stoll and Rudebeck, 2023) mean that subjects were likely actively tracking the probability of reward to make their choices. It is this active tracking and updating of reward probability that is reflected in the dynamics we observed at the neural population level in area 12o.

Anatomically, 12o is not only part of Carmichael and Price’s orbital network, as it has connections to other ventral frontal cortex subdivisions, but is also interconnected with areas within the medial frontal cortex (Carmichael and Price, 1996; Price, 2007). In particular, 12o has mono-synaptic connection with anterior cingulate cortex (ACC) (Jezzini et al., 2021; Trambaiolli et al., 2022), an area that is closely associated with guiding information seeking and exploratory behavior (Kolling et al., 2012, 2016; Stoll et al., 2016; Jezzini et al., 2021) and vlPFC, ACC, and amygdala are part of a network of strongly connected areas (Ghashghaei et al., 2007; Zeisler et al., 2023). If reward probability representations are a prominent feature of encoding in area 12o, it is possible that these representations regulate exploratory behavior by signaling the likelihood of receiving a reward when pursuing a specific stimulus in the current local environment or whether a new reward context should be sought out. Regarding whether such a role in reward learning and credit assignment extends to neighboring subdivisions (Monosov and Rushworth, 2022), our current work shows that neurons within 12o and 12l exhibited very different dynamics, indicative of functional specialization.

### The influence of cytoarchitecture and connectivity on ventral frontal representations

Even though our results identified clear distinctions in how cytoarchitecturally defined subdivisions of ventral prefrontal cortex represent information, it is unlikely that a given representation, or its associated function, entirely stop at a defined anatomical boundary. It is more likely the case that the precise patterns of inputs and outputs to the constituent neurons shape an area’s function, whereas the cytoarchitecture of an area constrains how that information is processed. For example, our study revealed that the two neighboring vlPFC areas 12o and 12l exhibited very different representations and dynamics. Despite this, connections between vlPFC and ACC span both 12o and 12l (Trambaiolli et al., 2022) and prior neuroimaging work reported that the representations of probability were present in both 12o and 12l subdivisions (Kaskan et al., 2017). Even though it is not possible to extract the full anatomical connectivity maps from animals undergoing neurophysiological recording, considering the anatomical connectivity of where neurons were recorded from is therefore critical when trying to understand the precise role of a given area. Only with this information in hand will it be possible to fully characterize how ventral frontal cortex contributes to reward-guided behaviors.

## CONFLICT OF INTEREST

None

## AUTHORS CONTRIBUTIONS

Conceptualization and Methodology: FMS, PHR; Investigation, Data curation, Analysis, Software and Visualization: FMS; Writing – Original Draft and Review/Editing: FMS, PHR; Funding Acquisition and Supervision: PHR.

## ACKNOWLEDGEMENTS

This work was supported by a National Institute of Mental Health BRAINS award to PHR (R01s MH110822; MH132064), a young investigator grant from the Brain and Behavior Foundation (NARSAD) to PHR, a Philippe Foundation award to FMS, seed funds from the Icahn School of Medicine at Mount Sinai to PHR. We thank Marques Love and Dr. Patrick Hof for their help in defining neuroanatomical boundaries.

## Notes

### Competing Interest Statement

The authors have declared no competing interest.

